# The projection basis determines the information ceiling for perturbation prediction

**DOI:** 10.64898/2026.07.07.737004

**Authors:** Simone Bianco

## Abstract

Recent benchmarks show that deep-learning models for perturbation prediction do not outperform simple baselines operating in principal-component (PCA) space. We explain this with an information-theoretic ceiling: for any orthonormal projection basis **Φ**, the squared correlation between prediction and truth is bounded by the variance the basis explains (*r*^2^ ≤ VE), so no model complexity can recover signal discarded at projection. On chemical perturbations (sciPlex3, LINCS L1000), the eigenbasis of a gene association network captures only 10–12% of drug-response variance and yields chance-level predictions, while PCA captures 90–99%. Graph wavelets built on the same network recover ∼88%, localising the drug signal in high-frequency modes that the standard low-pass eigenbasis discards. On CRISPRa genetic perturbations the ranking inverts: the network basis outperforms PCA across all dimensions tested. Controls on topology, null networks and data leakage confirm the effect is structural. The right basis depends on the perturbation modality: PCA captures the variance that drives chemical responses, the network basis captures the cascade structure that drives genetic ones, and bases that access the network’s full graph spectrum (such as graph wavelets) recover both from the same topology.

## Main

Deep-learning models for single-cell perturbation prediction, including GEARS ^1^, CPA ^2^, scGen ^3^ and several foundation models, were recently shown not to outperform simple linear baselines ^4^. Concurrent benchmarks reached similar conclusions ^5–7^: Viñas Torńe et al. showed that accounting for systematic variation eliminates apparent model advantages; Bendidi et al. found that PCA outperforms foundation models across perturbation tasks; and Csendes et al. showed that low perturbation-specific variance in standard benchmarks limits all models to near-mean-predictor performance. These results challenge the premise that architectural complexity improves perturbation prediction, but they do not explain why PCA-based representations succeed where biologically motivated alternatives fail.

We propose that, for drug perturbation prediction, the answer lies not in model capacity but in the projection basis: the coordinate system in which both complex, state-of-the-art models and simpler reference ones operate. Most methods, whether explicitly (PRESCIENT ^8^, CellOT ^9^) or implicitly (through encoder bottlenecks), project gene expression into a low-dimensional subspace before predicting perturbation effects. Principal component analysis (PCA) is the de facto standard for this step, yet its empirical dominance over biologically motivated alternatives has never been systematically explained.

We study this question using a potential-gradient dynamics framework inspired by the Waddington epigenetic landscape ^10^. In this view, cells occupy positions on a potential surface, and perturbations drive them along the landscape’s gradient toward new steady states. Waddington Potential Networks (WPN) ^8^ formalize this intuition by parameterizing cellular velocity as the gradient of a learned scalar potential, ***v*** = −∇*U*. This framework is well suited to isolating basis effects for two reasons. First, the gradient constraint restricts the model to conservative (irrotational) velocity fields, preventing it from learning arbitrary dynamics that could mask a poor projection; the quality of the basis is therefore directly reflected in prediction accuracy. Second, the projection basis enters as the model’s sole input representation: the network learns a scalar potential over projected coordinates, so changing the basis changes what the model can see without altering how it reasons about dynamics. We hold all other hyperparameters constant across experiments so that every observed difference in performance is attributable to the projection basis alone.

### An information-theoretic ceiling from orthogonal projection

Every method that reduces gene expression to a fixed low-dimensional coordinate system before predicting perturbation effects passes the signal through a linear bottleneck. For such methods, basis choice represents a hard cap on what any downstream model can recover.

### Proposition (projection bound)

For an orthonormal basis **Φ** ∈ ℝ*^G^*^×*k*^ and any predictor 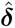 ∈ range(**Φ**), the squared Pearson correlation between truth and prediction, *r*^2^ := Corr(***δ***, 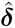)^2^, satisfies

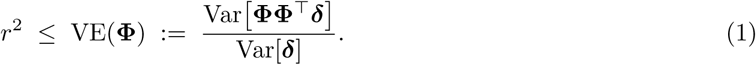

Decompose the true delta as ***δ*** = ***δ _‖_***+***δ***_⊥_ where ***δ _‖_***= **ΦΦ**^⊤^***δ*** lies in range(**Φ**). Because 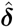 ∈ range(**Φ**) and ***δ***_⊥_ ⊥ range(**Φ**), we have Cov(***δ***_⊥_, 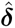) = 0, and Corr(***δ***, 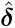)^2^ = Cov(***δ _‖_***, 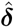)^2^*/*(Var(***δ***) Var(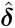)) ≤ Var(***δ _‖_***)*/* Var(***δ***) = VE(**Φ**) by Cauchy–Schwarz, with equality iff 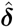 ∝ ***δ*** _||_.

The bound (1) is elementary. We state it explicitly because it separates two distinct bottlenecks that empirical work on perturbation prediction often conflates. Information availability is the variance explained VE(**Φ**), that is, the fraction of task-relevant signal the basis preserves. A basis with low VE cannot support high-accuracy predictions at any model complexity. Information exploitation is the gap *r*^2^*/*VE(**Φ**), or how much of the available signal a given architecture recovers. The bound cannot diagnose exploitation, but it upper-bounds what exploitation can achieve.

### Experimental framework

We test the ceiling through three complementary experiments chosen so that they can independently falsify or support the framework. Test 1 (chemical perturbations: sciPlex3 across three cell lines; LINCS L1000 cross-platform) asks whether a low-VE basis supports above-chance prediction in principle; Test 2 (graph wavelets on the same topology) asks whether restoring variance access to the same graph recovers the missing signal; Test 3 (Norman 2019 CRISPRa genetic perturbations) tests the modality-dependence prediction that graph-spectral bases should win on perturbations whose responses align with graph geometry. We employ a set of mechanistic controls (degree-preserving permutation, synthetic ground-truth network, PCA fit on untreated cells) to rule out confounders, including dimensionality artefacts, network miscalibration, and basis circularity.

### Data

We test our hypothesis on a number of prototypical perturbation response data sets across modalities. sciPlex3^11^ is a large-scale chemical perturbation dataset profiled with single-cell RNA sequencing across 188 drugs × 4 doses in three cell lines (K562, A549, MCF7; 8,288 genes, 2,202 conditions after filtering). LINCS L1000^12^ profiles 6,282 (MCF7) and 4,524 (A549) chemical perturbations on a bead-based 978-landmark gene platform. Norman et al. 2019^13^ profiles 236 single-gene CRISPRa activations in K562 (91,168 cells, 2,257-gene intersection with the network vocabulary). All experiments use per-cell-line controls; we identified and corrected a pooled-control artefact in sciPlex3 where cross-cell-line control pooling inflates apparent accuracy by allowing models to predict cell-line identity rather than drug effects (Appendix 1).

### Models

The Waddington Potential Network (WPN) projects gene expression onto a *k*-dimensional basis, learns a scalar potential via a residual Multi-Layer Perceptron (MLP; 19.6M parameters), and derives velocity as ***v*** = −∇***_s_****U* (***s***) via automatic differentiation. We additionally train a simple 3-layer residual MLP (79K–525K parameters, no gradient structure) and a scGen-style variational autoencoder on the same bases to separate basis effects from architectural ones. Predictions are evaluated on held-out conditions using direction accuracy (dir_acc), Pearson correlation on the top 200 differentially expressed genes (DEGs), and solver consistency (Online Methods).

### A low variance basis limits chemical perturbation predictions

Test 1 asks whether a low-VE basis can support above-chance prediction of chemical drug responses in principle. By Eq. (1), if the gene-association Laplacian eigenbasis captures only a small fraction of drug-delta variance, then *r* is capped at 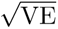 regardless of model complexity. We first probe this ceiling in three cell lines on the sciPlex3 dataset.

We constructed a composite gene-gene association network (534,670 edges) from protein-protein interactions (STRING ^14^), literature-curated signaling (OmniPath ^15^), and pathway co-membership edges (Reactome, KEGG, Gene Ontology; Online Methods). This network (hereafter GRN, following convention in the perturbation prediction literature) is primarily associative: its edges reflect functional co-occurrence and shared pathway annotation rather than directed regulatory causation (see Discussion). We computed the normalized Laplacian **L** = **I** − **D**^−1*/*2^**AD**^−1*/*2^ and extracted the lowest 256 non-trivial eigenvectors as a spectral basis.

This network eigenbasis explains only 11.8% of drug-response variance (measured on drug deltas ***δ*** = ***x̄***_treat_ − ***x̄***_ctrl_; Fig. 1a). In comparison, PCA recapitulates 90.1% of the original signal, while a random orthogonal basis reconstructs 3.0%. To confirm that this 11.8% reflects genuine biological topology rather than the degree distribution, we ran a degree-preserving permutation test: permuted networks with identical degree sequences capture only 4.2% ± 2.6% of variance (*p <* 10^−4^; Appendix 3). The network eigenvectors describe smooth, coordinated variation across co-regulated gene modules, patterns in which neighboring genes in the network change together. However, drug responses do not follow this structure: for example, a kinase inhibitor affects a sparse set of downstream targets scattered across unrelated network neighborhoods, producing gene-level changes that vary sharply between adjacent nodes in the graph ^16^. Such signals are high-frequency in the graph-spectral sense and lie outside the subspace spanned by the low-frequency eigenbasis. In physical terms, the network eigenbasis acts as a low-pass filter that captures only the smooth, coordinated modes of the association network, but drug responses behave more like impulse noise, activating scattered targets that cut across the network’s natural resonances.

**Figure 1:**
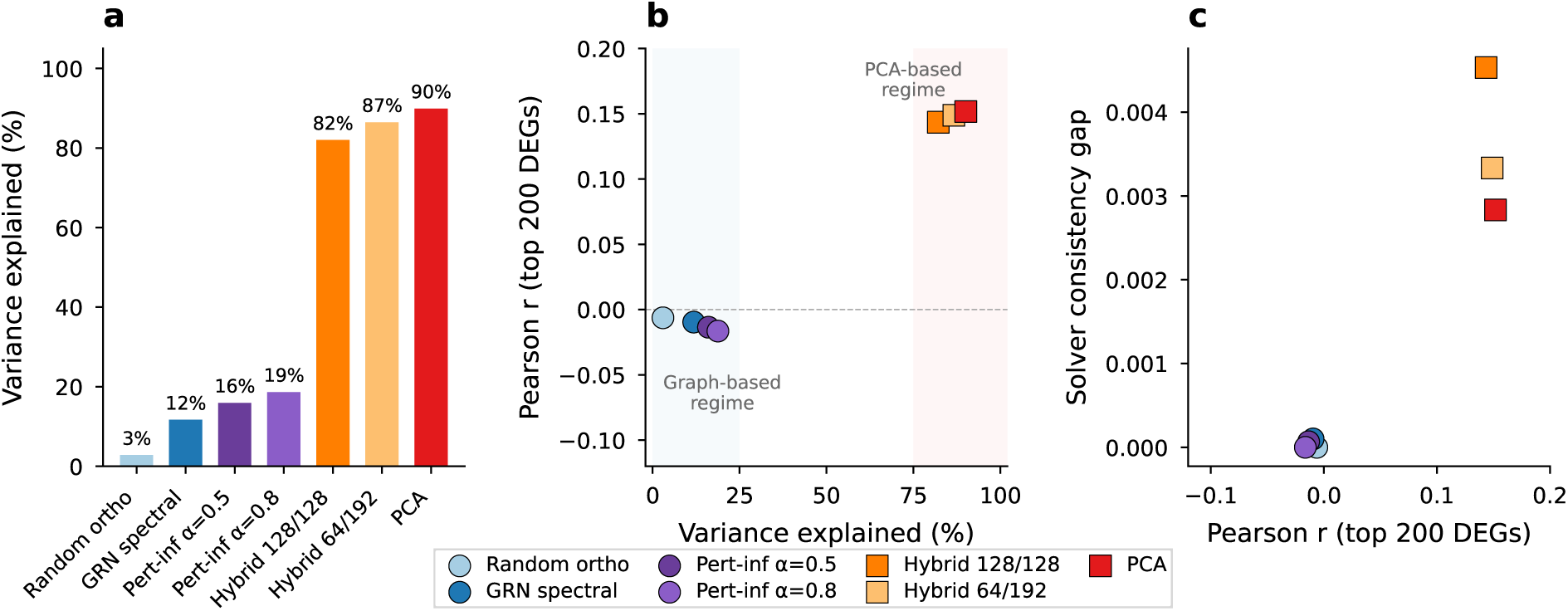
Projection basis determines the information ceiling for drug perturbation prediction. **a**, Fraction of drug-response variance (Var[**ΦΦ**^⊤^***δ***]*/*Var[***δ***]) captured by each 256-dimensional basis on K562 sciPlex3 drug deltas. GRN-spectral: 11.8%; perturbation-informed (*α*=0.8): 18.8%; PCA: 90.1%. **b**, Pearson correlation (*r*) on held-out conditions vs. variance explained. The tested bases cluster into two groups: graph-based (*<*20% variance, *r* ≈ 0) and PCA-containing (*>*80% variance, *r* ≈ 0.15), though the underlying relationship is continuous and monotonic (Supplementary Table S5). Dashed line indicates *r* = 0. **c**, Solver consistency gap vs. Pearson correlation. All gaps are small (*<*0.005), indicating stable dynamics across all bases.

To quantify this spectral mismatch, we computed the Graph Fourier Transform (GFT) of all drug deltas on the full 8,288-dimensional network Laplacian (Extended Data Fig. ED1). The mean power spectrum is nearly flat across all graph frequencies, with normalized spectral entropy *H/H*_max_ = 0.935 on K562, close to the maximum of 1.0 for a uniform distribution. This spectral diffuseness is not cell type-specific: A549 and MCF7 show similarly high entropy (0.933 and 0.890, respectively), confirming that drug responses are spectrally diffuse on the association network across all three cell lines. Cumulative energy curves confirm the disparity: PCA-256 captures ∼90% of drug-delta variance, while the network eigenbasis captures only ∼12% (Extended Data Fig. ED1b). This spectral diffuseness establishes that the network basis failure is biological, not circular: drug responses produce high-frequency signals on the association graph that low-frequency Laplacian eigenvectors cannot represent. Since PCA is fitted to training drug data, this comparison could appear circular; however, PCA computed solely on untreated control cells still captures 65–85% of drug-response variance, far exceeding the network eigenbasis (see Mechanistic controls).

### Replication across three cell lines

To test whether variance starvation is specific to K562, we repeated the GRN-spectral and PCA comparison on A549 (lung epithelial, 795 conditions) and MCF7 (breast epithelial, 656 conditions) using per-cell-line controls (Table 1). The pattern is universal and reproducible across random seeds (3 seeds each): GRN-spectral predictions are at or below chance (dir_acc = 0.43–0.49, *r* ≤ 0.06), while PCA achieves above-chance accuracy (0.58–0.66, *r* = 0.17–0.38). PCA captures a higher fraction of drug-delta variance in A549 (97.7%) and MCF7 (98.6%) than in K562 (90.1%), tracking the stronger PCA-based performance in those cell lines and reinforcing the VE→ *r* coupling predicted by Eq. (1).

**Table 1:** Cross-cell-line comparison of GRN-spectral vs. PCA on sciPlex3. All experiments use per-cell-line controls and the identical WPN architecture. All values are means across 3 random seeds; ± denotes standard deviation.

| Cell line | Basis | Var. expl. | dir_acc↑ | Pearson↑ | MSE<sub>20</sub>↓ |
| --- | --- | --- | --- | --- | --- |
| K562 | GRN-spectral | 11.8% | 0.433±.029 | 0.029±.049 | 1.100±.037 |
| | PCA | 90.1% | **0.602**±.003 | **0.168**±.017 | **0.137**±.007 |
| A549 | GRN-spectral | 10.3% | 0.490±.017 | 0.057±.013 | 7.08±.11 |
| | PCA | 97.7% | **0.656**±.009 | **0.384**±.043 | **1.11**±.15 |
| MCF7 | GRN-spectral | 10.1% | 0.482±.019 | -0.077±.040 | 37.3±.5 |
| | PCA | 98.6% | **0.583**±.003 | **0.220**±.014 | **2.77**±.49 |

### Unseen-drug generalization

To rule out drug-level memorization, we held out 20% of drugs (37 drugs, 148 conditions) from K562 and retrained both bases on the remaining 151 drugs (603 conditions). PCA generalizes to unseen compounds: dir_acc = 0.602 on test drugs vs. 0.601 on training drugs; Pearson *r* = 0.185 vs. 0.151 (Table 2). GRN-spectral remains at chance on both splits. We further confirm this with cross-cell-line PCA transfer: PCA fit on one cell line’s drug deltas captures 39–68% of variance on a different cell line, always far exceeding the network eigenbasis (see Mechanistic controls).

**Table 2:** Drug generalization on K562 sciPlex3. Models trained on 80% of drugs (151 drugs, 603 conditions) and evaluated on held-out drugs (37 drugs, 148 conditions). The accuracy gap between bases is preserved on unseen compounds.

| Basis | Split | dir_acc↑ | Pearson↑ | MSE<sub>20</sub>↓ | Gap↓ |
| --- | --- | --- | --- | --- | --- |
| GRN-spectral | Train drugs | 0.429 | 0.022 | 1.046 | 0.000 |
|  | Unseen drugs | 0.457 | 0.077 | 1.082 | 0.000 |
| PCA | Train drugs | 0.601 | 0.151 | 0.117 | 0.002 |
|  | Unseen drugs | **0.602** | **0.185** | 0.164 | 0.002 |

### Cross-platform replication (LINCS L1000)

The sciPlex3 results could in principle reflect an scRNA-seq-specific artefact. We tested the same PCA-vs-GRN comparison on LINCS L1000^12^, a bead-based 978-landmark platform with completely different measurement physics and orders of magnitude more conditions (MCF7: 6,282 conditions, 143K cells; A549: 4,524 conditions, 69K cells; 596-gene intersection with the STRING network). The variance gap between PCA and GRN is confirmed (Table 3, Supplementary Fig. S3): at *k*=256, PCA captures 95.8–97.0% of perturbation-delta variance vs. 48.4–49.3% for GRN, and the mean predictor’s Pearson *r* tracks VE faithfully (PCA *r* = 0.368–0.472 vs. GRN *r* = 0.268–0.347). The GRN basis reaches higher absolute VE on LINCS (∼49%) than on sciPlex3 (∼12%) because LINCS restricts to 596 landmark genes whose covariance is more concentrated along the network’s smooth modes; yet even in this easier regime, PCA-*k* at every *k* dominates GRN-*k* by an order of magnitude at small *k* and ∼2× at *k*=256. The basis effect is therefore not an artefact of scRNA-seq sparsity, drug panel, or gene set.

**Table 3:** LINCS L1000 cross-platform replication. The PCA ≫ GRN variance gap reproduces on a bead-based measurement platform with a different gene panel (596-gene intersection with STRING), different cell populations, and orders of magnitude more conditions. Mean-predictor Pearson *r* tracks variance explained as predicted by Eq. (1), across both cell lines.

| Cell line | $k$ | PCA VE | GRN VE | Mean-PCA $r$ | Mean-GRN $r$ |
| --- | --- | --- | --- | --- | --- |
| MCF7 | 10 | 72.7% | 8.7% | 0.333 | -0.003 |
| MCF7 | 50 | 88.0% | 16.1% | 0.363 | 0.101 |
| MCF7 | 256 | 97.0% | 48.4% | 0.368 | 0.268 |
| A549 | 10 | 70.4% | 10.4% | 0.402 | 0.091 |
| A549 | 50 | 84.0% | 17.6% | 0.460 | 0.157 |
| A549 | 256 | 95.8% | 49.3% | 0.472 | 0.347 |

Test 1 therefore corroborates the availability bottleneck across three cell lines, two measurement platforms, and held-out drugs: chemical drug responses are spectrally diffuse on gene-association graphs, and the low-pass eigenbasis starves the downstream model of signal that no architecture can recover.

### Basis dependence is architecture-independent

To confirm that the availability ceiling is a property of the basis rather than of our particular model, we replicated the sciPlex3 K562 comparison across three architectures and seven bases. These include pure PCA, the GRN low-pass eigenbasis, a random orthogonal basis, two perturbation-informed reweightings of the GRN (*α* ∈ {0.5, 0.8}), and two Hybrid-*m*/*n* bases that concatenate the top *m* GRN eigenvectors with the top *n* = 256−*m* PCA components and re-orthogonalise the result by QR decomposition (Online Methods; Table 4, Fig. 1). The bases cluster into two regimes separated by their VE on drug deltas, though the underlying VE-vs-performance relation is continuous (Supplementary Table S5). Graph-based bases (GRN-spectral, random orthogonal, perturbation-informed; VE ≤ 19%) all produce predictions at or below chance, with near-zero Pearson values. PCA-based bases (Hybrid-128/128, Hybrid-64/192, pure PCA; VE ≥ 82%) all achieve consistent above-chance performance, with PCA highly stable across seeds (±0.003 dir_acc). The regime-boundary effect reaches calibration as well: low-variance bases produce MSE_20_ nearly 10× above PCA-based models (1.0–1.3 vs. 0.13–0.14) and magnitude ratios of 3.9–7.7× (Table 1), consistent with overfitting to noise dimensions in variance-poor subspaces. To verify that the transition is monotonic and entirely determined by VE, we ran a PCA dimensionality sweep over *k* = 10–512 (Extended Data Fig. ED2).

**Table 4:** Basis and model comparison on K562 sciPlex3 (256 dimensions, per-cell-line controls). All WPN rows use the identical architecture (19.6M parameters, 40 epochs). Linear baselines operate in the same projection spaces. Var. expl. measured on drug deltas. Chance-level dir_acc = 0.50. ± values: standard deviation across 3 random seeds.

| Model / Basis | Var. expl. | Params | dir_acc↑ | Pearson↑ | MSE<sub>20</sub>↓ |
| --- | --- | --- | --- | --- | --- |
| Linear baselines |  |  |  |  |  |
| Mean predictor (gene space) | 100% | 0 | **0.650** | **0.216** | 0.134 |
| Ridge (PCA-256, dose) | 90.1% | 512 | 0.621 | 0.175 | 0.134 |
| Mean (GRN-256 projected) | 11.8% | 0 | 0.600 | 0.050 | 0.143 |
| WPN, graph-based (<20% variance) |  |  |  |  |  |
| Random orthogonal | 3.0% | 19.6M | 0.402 | -0.006 | 1.291 |
| GRN-spectral | 11.8% | 19.6M | 0.433±.029 | 0.029±.049 | 1.100±.037 |
| Pert-informed ($\alpha=0.5$) | 16.1% | 19.6M | 0.428 | -0.013 | 1.074 |
| Pert-informed ($\alpha=0.8$) | 18.8% | 19.6M | **0.436** | -0.016 | **1.032** |
| WPN, PCA-based (>80% variance) |  |  |  |  |  |
| Hybrid-128/128 (GRN + PCA) | 82.2% | 19.6M | 0.588 | 0.144 | 0.136 |
| Hybrid-64/192 (GRN + PCA) | 86.6% | 19.6M | 0.600 | 0.149 | 0.135 |
| PCA | 90.1% | 19.6M | 0.602±.003 | 0.168±.017 | 0.137±.007 |

### The dichotomy is architecture-independent on chemical perturbations

A 3-layer residual MLP (525K parameters) reproduces the same dichotomy on K562 sciPlex3: GRN-spectral = 0.418 ± 0.004, PCA = 0.610 ± 0.002 (3 seeds; Extended Data Table ED1), values that match the WPN’s 0.433 ± 0.029 and 0.602 ± 0.003 despite 37× fewer parameters and no gradient-based dynamics. An scGen-style variational autoencoder reproduces the same pattern (Supplementary Table S10).

### Linear baselines match the parameterised models

In PCA-256 space on K562 sciPlex3, the zero-parameter mean predictor achieves dir_acc = 0.650, *r* = 0.216, slightly exceeding the WPN (0.602, 0.168), and a Ridge regression with dose as its sole feature achieves dir_acc = 0.621 and *r* = 0.175. This is expected given sciPlex3’s shared-control design: all K562 conditions have the same control mean, so the WPN computes a single velocity ***v*** = −∇*U* (***s***_ctrl_) for all conditions, and the MSE-optimal single vector is the mean delta. The potential-gradient architecture cannot escape the ceiling when the input provides no condition-specific information; what distinguishes the projected from the gene-space mean predictor is solely the amount of variance preserved by the basis. The same mean predictor projected through the network basis drops from *r* = 0.216 to *r* = 0.050, again confirming that basis choice rather than architecture sets the ceiling.

### Hybrid bases track variance, not biology

The Hybrid-*m*/*n* bases produce performance that tracks PCA VE in strict proportion: Hybrid-128/128 (82.2% VE) → *r* = 0.144; Hybrid-64/192 (86.6%) → *r* = 0.149; pure PCA-256 (90.1%) → *r* = 0.168 (Fig. 1b,c). The network components contribute negligibly, providing neither additional prediction signal nor measurable regularisation (solver consistency gaps 0.003–0.005; Fig. 1c). The network eigenvectors in these bases are effectively inert dimensions that reduce variance coverage relative to pure PCA without offering compensating advantages.

### Edge reweighting and directed regulatory networks do not rescue the eigenbasis

A natural rescue strategy is to keep the topology but retune the edge weights to drug-response relevance. We therefore tested perturbation-informed reweighting, where each edge weight is interpolated between the composite topology confidence and the absolute Pearson correlation |*ρ_ij_*| of genes *i* and *j* across all drug-condition deltas: 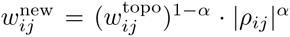, with *α* ∈ [0, 1] (Online Methods). A full sweep from *α* = 0 to *α* = 1 (Extended Data Fig. ED4) lifts VE from 11.8% to 18.8% at best (*α* = 0.8) but never produces above-chance prediction (dir_acc = 0.436, *r* = −0.016; Table 4). Network sparsification at confidence thresholds between 0.1 and 0.9 also fails to lift VE above 20% (Extended Data Fig. ED3), and a cell-type-specific network built from K562 control-cell gene–gene correlations peaks at 21% VE (Supplementary Table S6). Stratifying the 188 sciPlex3 drugs into 18 mechanism-of-action classes confirms that no drug class is preferentially aligned with the GRN basis (Supplementary Table S7). Replacing the undirected composite graph with curated directed TF–target databases (CollecTRI ^17^, DoRothEA ^18^) also fails: these networks cover fewer than half of the 8,288 genes and produce lower VE (4.7–6.2%; Supplementary Table S12).

### Wavelet rescue and spectral localisation on the same topology

Test 2 asks the converse of Test 1: if chemical drug responses are spectrally diffuse on the association graph, can we recover the missing information by enlarging the basis to access the same graph’s higher frequencies? If wavelet bases that span the full Laplacian spectrum rescue prediction, the failure in Test 1 is specifically a low-pass failure, not a network-topology failure.

We constructed three graph-wavelet bases that access all network frequencies: band-pass filtering (partitioning the Laplacian spectrum into *J* =8 frequency bands and selecting the most informative directions within each), spectral graph wavelets ^19^ (*g*(*x*) = *xe*^−^*^x^* at 8 scales), and diffusion wavelets (*e*^−^*^tL^* at 8 timescales; Supplementary Table S4). Graph wavelets have recently been applied to single-cell data for gene embedding ^20^; here we use them as projection bases for perturbation prediction.

### Wavelet bases match PCA on the same topology

Band-pass and diffusion wavelets recover nearly 90% of drug-delta variance on K562, on par with PCA in the same experiment (band-pass VE = 88.1%; diffusion VE = 88.8%; PCA reference VE = 89.4%), while the standard low-pass network eigenbasis captures only 11.8%. The mean predictor in wavelet-projected space achieves *r* = 0.211, comparable to the gene-space mean (*r* = 0.216). A trained MLP on band-pass and diffusion wavelets achieves *r* = 0.155–0.156, matching PCA-MLP (*r* = 0.153); the spectral wavelet MLP is lower (*r* = 0.093), consistent with its lower variance coverage (82% vs. 88–89%). The same pattern replicates on A549 and MCF7, where bandpass and diffusion wavelets recover 97–98% of drug-delta variance versus 10% for the low-pass eigenbasis (Supplementary Table S4).

### Drug-response signal is localised in a high-frequency band

To localise where in the Laplacian spectrum the drug-response signal lives, we partitioned the spectrum into *J* =8 logarithmic bands and measured the fraction of drug-delta variance carried by each band. On K562 sciPlex3, band 7 (*λ* ∈ [0.326, 2.0], containing 8,235 eigenvectors) accounts for 91.9% of total drug-delta variance, while the lowest band (*λ* ∈ [0, 0.02], containing the 256 smoothest eigenvectors used by the standard network eigenbasis) contributes *<*10% (Supplementary Fig. S4). The low-pass eigenbasis and the drug-response signal therefore live in almost disjoint subspaces of the same Laplacian, suggesting that the signal is not reachable. This establishes that the failure of the network eigenbasis in Test 1 is specifically a spectral-range failure, not a topology failure: VE can be raised from 12% to 88% on the identical graph by replacing the low-pass filter with band-pass or diffusion wavelets. Moreover, this confirms the results of Test 1: Laplacian eigenvectors minimise Dirichlet energy and are therefore the smoothest possible basis on any graph, regardless of edge weights ^16^, so reweighting cannot bypass the low-pass restriction that the wavelet analysis shows is the binding constraint.

Graph wavelets thus demonstrate that the network topology contains the drug-response information Test 1 showed the eigenbasis lacks, and they offer a path that incorporates biological structure without sacrificing variance coverage.

### Genetic perturbations invert the ranking

Tests 1 and 2 establish that chemical perturbations are spectrally diffuse on the association graph. Our Test 3 attempts at addressing whether this failure is a property of the basis or of the perturbation modality. Unlike pharmacological inhibitors, which hit sparse and scattered molecular targets, single-gene CRISPR activations propagate along regulatory cascades that follow network edges. For genetic perturbations, this would mean that the low-frequency modes the network eigenbasis captures should carry more of the signal, and parameterised models on the network basis should outperform those on PCA.

We tested this prediction on Norman et al. 2019^13^, a CRISPRa activation dataset (91,168 cells, 236 single-gene activations in K562), restricted to the 2,257-gene intersection with the network vocabulary (54,844 edges). We held *n*_train_ = 189 perturbations and evaluated at basis dimensionalities *k* ∈ {25, 50, 100, 150, 189} using three random seeds.

### Mean predictor: VE still wins

The zero-parameter mean predictor in gene/PCA space achieves dir_acc = 0.887 ± 0.023, *r* = 0.648 ± 0.017 (3 seeds), exceeding the network-projected mean (dir_acc = 0.805 ± 0.022, *r* = 0.370 ± 0.012; Table 5, Supplementary Fig. S1). Consistent with sciPlex3, PCA preserves substantially more variance (100% vs. 19.2%), and the availability gap persists: the basis ceiling governs the mean predictor on both modalities.

**Table 5:** Test 3, Genetic perturbation inversion on Norman 2019 CRISPRa (K562, 236 single-gene activations, 2,257-gene network intersection, *k* = 189, 3 seeds). The mean predictor still favours PCA (availability ceiling preserved), but both parameterised architectures invert the ranking and favour the network basis, the opposite of Table 4. Full *k*-sweep in Supplementary Table S2.

| Model | Basis | Var. expl. | Params | dir_acc↑ | Pearson↑ |
| --- | --- | --- | --- | --- | --- |
| Mean predictor baselines |  |  |  |  |  |
| Gene space | — | 100% | 0 | **0.887**±.023 | **0.648**±.017 |
| PCA-189 | PCA | 100% | 0 | **0.887**±.023 | **0.648**±.017 |
| GRN-189 | GRN | 19.2% | 0 | 0.805±.022 | 0.370±.012 |
| Parameterised models (ranking inverts) |  |  |  |  |  |
| WPN | GRN-spectral | 19.2% | 15.8M | **0.723**±.013 | **0.143**±.010 |
| WPN | PCA | 100% | 15.8M | 0.587±.008 | 0.010±.003 |
| MLP | GRN-spectral | 19.2% | 163K | **0.725**±.016 | **0.142**±.009 |
| MLP | PCA | 100% | 163K | 0.591±.008 | 0.009±.005 |

### Parameterised models invert the ranking

The WPN trained on the network basis achieves dir_acc = 0.723 ± 0.013, *r* = 0.143 ± 0.010, outperforming the PCA-basis WPN (dir_acc = 0.587 ± 0.008, *r* = 0.010 ± 0.003) and exactly inverting the sciPlex3 ranking. A residual MLP (79K–163K parameters) on the same bases reproduces the inversion almost exactly at every *k* from 25 to 189 (Supplementary Table S3), ruling out a *k* = *n*_train_ overfitting artefact or a WPN-specific inductive bias. At *k* = 25, for instance, PCA still captures 87.3% of training-delta variance but its direction accuracy is 0.603 vs. 0.714 for GRN-25.

### Why the inversion is consistent with the spectral interpretation

On the learned potential landscapes at *k* = 189, both GRN-WPN and PCA-WPN converge to effectively rank-1 velocity fields, but the GRN-WPN’s velocity is substantially better aligned with the true deltas (median cosine similarity = 0.917 vs. 0.785; mean 0.758 vs. 0.668), with better-calibrated magnitudes (1.03 ± 0.71 vs. 0.90 ± 0.64). To rule out an optimisation-smoothness account of the inversion, we computed the condition number *κ* of the Hessian of the learned potential *U* at the control state ***s***_ctrl_, defined as the ratio of its largest to smallest eigenvalue; lower *κ* corresponds to a more isotropic, better-conditioned loss surface that is easier to optimise. PCA-WPN has the lower value (*κ* = 373 vs. 1,013 for GRN-WPN), so, although PCA is provided with the easier optimisation landscape, it still loses to GRN. This suggests that inversion is a geometry effect. The network basis restricts the model’s gradient to a subspace aligned with gene-association structure, which for CRISPRa perturbations (whose effects propagate along pathway edges) acts as an informative inductive bias. PCA, by contrast, provides *k* roughly equally weighted dimensions with no directional prior. Parameterised models therefore exploit basis geometry in ways that variance coverage alone does not predict, particularly on small datasets.

Test 3 therefore supports a refined version of the ceiling framework. The availability bottleneck (mean predictor VE) transfers from chemical to genetic perturbations without modification; what changes is which basis geometry is productively exploitable by parameterised models. The fact that the same information framework predicts the ranking on drugs and its inversion on CRISPRa strengthens our central claim that basis choice is a modality-dependent design decision with a provable ceiling.

### Mechanistic controls

To rule out network-miscalibration, basis-circularity, and dimensionality-artefact accounts of the Test 1–3 pattern, we ran three complementary controls (Supplementary Fig. S5).

### Control 1: Degree-preserving permutation null

If the 11.8% VE captured by the association network were simply an artefact of its heavy-tailed degree distribution, any network with the same degree sequence would perform equivalently. We generated 10 degree-preserving edge-swap rewirings ^21^ of the association graph (∼77% swap acceptance rate) and evaluated each as a spectral basis. The real network captures 11.8% of drug-delta variance vs. 4.2% ± 2.6% for the permuted null (*p <* 10^-4^ by one-sided permutation test; direction accuracy 0.601 vs. 0.544 ± 0.017, *p <* 10^-4^; Supplementary Table S11). The association topology therefore carries genuine biological information, but this information is not enough to bridge the gap to PCA.

### Control 2: Synthetic ground-truth network

As a positive control, we simulated a scale-free-like regulatory network (100 genes, 20 transcription factors, 103 directed edges, Hill-function dynamics), generated 500 control cells and 30 single-gene knockouts, and evaluated the true generative network, degree-preserving permutations, and random density-matched networks as eigenbases. The true network captures 44.2% of variance vs. 31.2 ± 1.5% (permuted) and 29.3 ± 2.2% (random; *p <* 10^−4^ for both comparisons; Supplementary Table S14). The permutation framework therefore detects genuine network structure when the data really are generated by it. Critically, even with perfect knowledge of the generative graph, the eigenbasis still captures less than half the variance that PCA achieves (∼100%), suggesting that, indeed, the low-pass smoothness constraint is limiting.

### Control 3: PCA on untreated controls breaks circularity

PCA fit on training drug deltas and then evaluated on the same deltas could appear circular: PCA is defined to maximise exactly the signal being evaluated. To break this link, we computed PCA on the raw control-cell expression matrix (cells that have never seen a drug) for each cell line independently and measured variance explained on drug deltas. This “control-PCA” basis captures **65–85%** of drug-response variance across all three sciPlex3 cell lines, compared to 10–12% for the network eigenbasis, and its mean-predictor Pearson is virtually identical to drug-fit PCA (K562: 0.163 vs. 0.163; A549: 0.342 vs. 0.343; MCF7: 0.236 vs. 0.237; Supplementary Table S15). The PCA advantage therefore reflects basal gene covariance structure, not circularity from including drug-response data in the basis computation.

## Discussion

In this work we have performed three tests to provide clarity between two distinct hypotheses behind the recent observation of deep perturbation-prediction models not beating PCA-based baselines. The first account attributes the gap to insufficient model capacity or inductive bias. The second, which we advance here, attributes it to an information-theoretic ceiling set by the projection basis. Our tests rule out the first account and support the second across three independent axes: Test 1 shows that chemical drug responses are spectrally diffuse on a gene-association graph (GRN), reducing the variance available to low-pass eigenbases, and the PCA ≫ GRN gap replicates across three cell lines and an independent measurement platform (LINCS L1000); Test 2 shows that the failure is specifically a spectral-range failure: band-pass and diffusion wavelets on the same topology recover nearly 90% of variance, and 91.9% of drug-delta variance lives in a disjoint high-frequency band that low-pass eigenbases exclude by construction; Test 3 shows that the ranking inverts on genetic perturbations exactly as the spectral interpretation predicts (CRISPRa responses propagate along regulatory cascades and align with low-frequency pathway-coherent modes), so parameterised models on GRN outperform those on PCA at every *k* tested, without any change to the underlying VE ceiling. Three mechanistic controls (degree-permutation null, synthetic ground-truth network, control-only PCA) close the loop by ruling out degree-distribution, network-miscalibration, and basis-circularity confounds.

### Availability vs. exploitation

The *r*^2^ ≤ VE bound separates two bottlenecks that prior empirical work often conflated. Availability (VE) is what Tests 1 and 3’s mean predictors track: PCA at 90% → *r* ≈ 0.22 ceiling on sciPlex3; GRN at 12% → *r* ≤ 0.35 ceiling; both behave exactly as Eq. (1) predicts. Exploitation (*r*^2^*/*VE) is what parameterised models add on top: in our data *r*^2^*/*VE *<* 15% across all 72 triples (Supplementary Fig. S2), and on sciPlex3’s shared-control design the potential-gradient inductive bias is actively costly relative to the mean. Practically, when condition-specific inputs are missing (sciPlex3’s pooled-control degeneracy), the MSE-optimal predictor is the unconditional mean, and projecting through any basis only subtracts information. The framework therefore gives a clean decomposition: VE is a property of the basis, and *r*^2^*/*VE is a property of the model × task × sample-size interaction. Both matter, but only the first is strictly fixed by the projection choice.

### Modality-dependence of exploitation

Test 3’s inversion shows that the ceiling framework accommodates modality-dependent basis choices. Pharmacological drugs hit sparse, scattered molecular targets and produce signals that are spectrally diffuse on any coherence-based graph; the low-frequency modes cannot represent them. CRISPR activations instead propagate along regulatory edges, so the low-frequency pathway-coherent modes contain most of the response, and the GRN basis provides an informative directional prior for parameterised models. The same spectral reading of Eq. (1) predicts both observations, and the inversion therefore confirms the central claim. Additionally, we perform a CRISPRi vs. CRISPRa dissociation test (Appendix 3) to show that the association network does not encode directional regulatory information sufficient to differentially aid one modality. Thus, the Test 3 inductive-bias benefit is plausibly a cascade-propagation unrelated to any signed-regulation effect.

### What does and does not follow for method design

Our framework predicts that any method passing through a fixed linear bottleneck of *k* dimensions is capped by VE*_k_* of that bottleneck; this includes scGen, CPA, chemCPA, and latent-space autoencoders that operate in ∼128-dim subspaces. Full-gene methods (GEARS ^1^, PRiMeFlow ^22^) are not directly constrained by the linear ceiling, but have nevertheless not outperformed PCA-based baselines on existing benchmarks ^4^, suggesting that exploitation (not availability) is their binding constraint at current sample sizes. Taken together, these observations allow the formulation of three practical guidelines. (i) Benchmarks should report VE of each method’s internal representation on task-relevant variation (drug or perturbation deltas, not total expression), not only the final prediction metric. (ii) Biologically motivated projections should be compared to degree-preserving permutation nulls, not just random projections (our Test 1 control). (iii) Graph wavelets are not the only remedy: any network-aware basis that spans the Laplacian’s full frequency range (rather than truncating to its low-pass head) recovers the variance the standard eigenbasis discards. The class includes the band-pass, spectral, and diffusion wavelets we test here, polynomial graph filters (Defferrard et al. ^23^, ChebNet-style), heat-kernel signatures at multiple scales, and any random projection drawn from the full eigenvector set. The Test 2 result therefore suggests that network topology, filtered with sufficient spectral coverage, is neither inert nor uninformative, and the choice between wavelet families is secondary to the choice not to truncate.

### Limitations

The primary tests are replicated across 3 seeds on sciPlex3 (K562, A549, MCF7) with both WPN and MLP architectures and on Norman 2019 with both WPN and MLP; single-run experiments (the *α* sweep, sparsity ablation, wavelet, scGen, hybrid bases) must be interpreted in proportion to effect size with some caution. The ceiling framework constrains methods with fixed linear bottlenecks; it does not directly bound nonlinear or full-gene architectures, and we do not benchmark GEARS, CPA, or PRiMeFlow. PerturBench-style rank metrics ^24^ (Supplementary Table S9) collapse to chance for all condition-invariant predictors under sciPlex3’s shared-control design, as expected: these metrics probe exploitation in settings with richer conditional structure than sciPlex3 provides. A more demanding exploitation test would use dose-response or combinatorial-perturbation datasets where the unconditional mean is no longer optimal. Re-evaluating on all 8,288 genes rather than the top-200 DEGs preserves the PCA ≫ GRN ordering (Supplementary Table S8), so the basis effect is not an artefact of the DEG-selection metric.

Linear-baseline benchmarks ^4,6,7^ have documented that PCA-based methods win on chemical perturbation tasks. This paper explains why, with three mutually reinforcing falsification tests, and it shows that the same framework predicts where PCA-based methods should lose: on genetic perturbations whose responses align with graph geometry. Basis choice is therefore a modality-dependent design decision with a provable ceiling, not a universally solved problem; and graph wavelets offer a concrete path that preserves both regimes.

The availability-versus-exploitation decomposition changes what a fair perturbation-prediction benchmark should measure. Reporting VE of each method’s internal representation alongside accuracy would drive the distinction of models that exploit their basis well from models that merely inherit a good projection. Experiments on datasets that break sciPlex3’s shared-control degeneracy (dose–response curves, combinatorial perturbations, and cross-modality transfer), where exploitation begins to matter and full-gene or mode-selective architectures may express advantages that linear bottlenecks forbid, should be clear next steps to establish a more rigorous framework for perturbation response benchmarking. We also predict that this framework applies to image-based perturbation readouts (Cell Painting, subcellular morphology), to spatial transcriptomics, and to any setting in which a fixed linear projection precedes a learned predictor, although proving this statement is outside the scope of this work.

## Code and data availability

All the code is available as attached software. All data used in this work is publicly available from the cited sources.

## Acknowledgements

The author would like to acknowledge critical discussions with Bikash Sabata, William Poole, Zichao Yan, Mehrshad Sadria and Esther Wershof.

## Author contributions

S.B. conceived the study, developed the framework, performed all analyses, and wrote the manuscript.

## Online Methods

### Data processing

sciPlex3^11^ raw UMI counts (GEO accession GSE139944) were aligned to the 8,288-gene vocabulary of the Tahoe-100M corpus ^25^ by matching gene symbols; genes absent from the vocabulary were discarded. For each drug-dose-cell-line condition (*n* ≥ 20 cells), we computed the mean expression vector. Per-cell-line control means were computed from control cells stratified by cell line metadata (K562: 3,935 cells; A549: 5,857; MCF7: 7,786). Drug deltas are defined as ***δ*** = ***x̄***_treat_ − ***x̄***_ctrl_ where ***x̄***_ctrl_ is the cell-line-specific control mean. The primary ablation uses K562 (751 conditions, 188 drugs × 4 doses) with an 80/20 train/validation split by condition. Cross-cell-line experiments use A549 (795 conditions) and MCF7 (656 conditions) with the same procedure. For the drug generalization experiment (Table 2), the split is by drug identity: 151 drugs (603 conditions) for training, 37 drugs (148 conditions) for testing, ensuring no drug-level leakage between splits. Multi-seed experiments repeat the K562 ablation with 3 independent random seeds (42, 1, 2) for data splitting and report mean ± standard deviation.

#### Norman 2019 CRISPRa

The preprocessed Norman et al. 2019^13^ dataset (91,168 cells, 5,575 genes, 236 single-gene CRISPRa activations in K562) was restricted to the 2,257-gene intersection with the network vocabulary. Mean expression vectors were computed per perturbation condition (*n* ≥ 20 cells) and split 80/20 by perturbation identity (189 train, 47 validation). PCA dimensionality was capped at *k* = 189 = *n*_train_ to avoid rank deficiency.

### Gene-gene association network

We constructed a composite gene-gene association network from STRING v12^14^ (protein-protein interactions, score ≥ 700) and OmniPath ^15^ (literature-curated signaling), supplemented with Reactome, KEGG, and Gene Ontology biological process co-annotation edges. Edges were filtered to confidence ≥ 0.1, yielding 534,670 edges over 8,288 genes. The adjacency matrix was symmetrized (directed edges made bidirectional).

### Projection bases

#### GRN-spectral

The normalized graph Laplacian **L** = **I**−**D**^−1*/*2^**AD**^−1*/*2^ was eigendecomposed via `torch.linalg.eigh`. The lowest *k*=256 non-trivial eigenvectors form **Φ**_GRN_ ∈ R*^n^*^×^*^k^*.

#### PCA

Computed on training data only (control mean + treated condition means). The top *k* right singular vectors form **Φ**_PCA_.

#### Random orthogonal

QR decomposition of a random Gaussian matrix (seed 42).

#### Hybrid

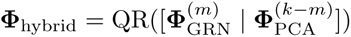, where superscripts denote the number of columns retained. QR preserves the network columns exactly and adjusts PCA directions to be orthogonal to the network subspace.

#### Perturbation-informed

For each edge (*i, j*) in the network, we compute |*ρ_ij_*| = |corr(***δ***_:*,i*_*, **δ***_:*,j*_)| across all training drug conditions. Edge weights are set to 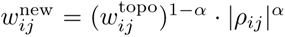. Edges with 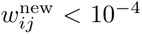 are removed. The Laplacian is recomputed and eigendecomposed.

#### Graph wavelets

We constructed three wavelet bases from the network Laplacian, each accessing the full frequency spectrum rather than only low-frequency modes. Band-pass: the Laplacian eigenvalues are partitioned into *J* =8 equal-width frequency bands, and within each band the eigenvectors explaining the most drug-delta variance are selected to fill 256 total dimensions. Spectral graph wavelets ^19^: the generating kernel *g*(*x*) = *x e*^−^*^x^* is evaluated at 8 logarithmically spaced scales, yielding 8 × *n* wavelet coefficients from which we retain the 256 highest-variance directions. Diffusion wavelets: the matrix exponential *e*^−^*^tL^* is computed at 8 timescales; the 256 highest-variance columns are retained. All three bases are orthonormalized via QR decomposition.

#### Cell-type-specific network

We constructed an alternative network from gene–gene Pearson correlations computed on K562 control cells (*n* = 3,935). Edges with |*r*| *< τ* are removed (thresholds *τ* ∈ {0.05, 0.1, 0.2, 0.3, 0.5}). The resulting adjacency matrix is symmetrized and processed identically to the STRING-based network (Laplacian eigendecomposition, lowest *k*=256 eigenvectors).

#### RandPCA

*N* columns are drawn uniformly at random (without replacement) from the full set of PCA eigenvectors, deliberately omitting the top components. This produces bases with low variance explained despite using PCA directions, serving as a control for the VE–performance relationship (Supplementary Table S5).

### Waddington Potential Network (WPN)

The model learns a scalar potential *U* : R*^k^* → R parameterized as a 6-layer residual MLP with SiLU activations, LayerNorm, and hidden dimension 1,024. Additional components include multi-scale potentials (*U*_fast_, *U*_slow_; 3-layer MLPs, hidden dim 512) and a perturbation-conditioned deformation Δ*U* (***s***; ***a***) (4-layer FiLM-conditioned MLP ^26^). The cellular velocity is ***v*** = −∇***_s_****U* (***s***), computed exactly via torch.autograd.grad. Total: 19.6M parameters.

Predicted gene expression is obtained by projecting to spectral space, one-step Euler integration (***ŝ*** = ***s***_ctrl_ + ***v***), back-projection, and residual refinement via a bottleneck MLP.

### Training

We train all models using the AdamW optimizer with a learning rate of 3 × 10^−4^ and weight decay 10^−4^, applying cosine annealing over 40 epochs with a batch size of 32 and gradient clipping at norm 1.0. The loss function combines three terms: a gradient-matching loss ℒ = ||***v***_pred_ − (***s***_treat_ − ***s***_ctrl_)|| ^2^ that drives the predicted velocity toward the observed perturbation delta, an attractor stability penalty (*λ_s_*= 0.1) that encourages the control state to sit near a potential minimum, and a curvature regularization term (*λ_c_*= 0.01) estimated via the Hutchinson trace estimator ^27^ that prevents overly sharp potential landscapes.

### Evaluation metrics

All metrics computed on top 200 DEGs ranked by absolute true change: direction accuracy (fraction with correct sign); Pearson correlation; MSE_20_ (mean squared error on top 20 DEGs); solver gap (|dir_acc_1-step_ − dir_acc_2×0.5-step_|).

For the optimisation-landscape diagnostic in the Norman analysis, we additionally compute the Hessian of the learned potential at the control state, 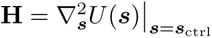, via `torch.autograd.functional.hessian`, and report its condition number *κ*(**H**) = *λ*_max_(**H**)*/λ*_min_(**H**) (eigenvalues of **H**, computed with `torch.linalg.eigvalsh`).

### Variance explained

Defined as VE = Σ*_j_* Var[(**ΦΦ**^⊤^***δ***)*_j_*] */* Σ*_j_* Var[***δ****_j_*], where the sum is over genes *j* and variance is computed across drug conditions. This measures the fraction of drug-response signal preserved by the project–back-project cycle.

#### Proposition (projection bound)

For an orthonormal basis **Φ** ∈ R*^G^*^×*k*^ and any predictor 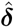 ∈ range(**Φ**), the squared Pearson correlation between truth and prediction (*r*^2^ := Corr(***δ***, 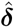)^2^) satisfies

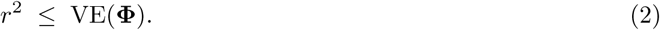

Proof. Decompose the true delta as ***δ*** = ***δ***_||_+ ***δ***_⊥_ where ***δ***_||_= **ΦΦ**^⊤^***δ*** lies in range(**Φ**) and ***δ***_⊥_ is orthogonal to it. Since 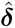 ∈ range(**Φ**), we have Cov(***δ***_⊥_, 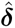) = 0. Therefore Corr(***δ***, 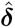)^2^ = Cov(***δ*** _||_, 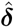)^2^*/*(Var(***δ***) Var(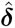)) ≤ Var(***δ***_||_)*/* Var(***δ***) = VE by the Cauchy–Schwarz inequality, with equality iff 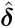 = *c **δ******_‖_*** for some scalar *c >* 0. This is a standard consequence of orthogonal projection geometry; we state it explicitly because it clarifies the role of variance explained as a necessary (though not sufficient) condition for prediction accuracy. All 72 data points in this study satisfy the bound (Supplementary Fig. S2). The more revealing observation is how far below the bound all points fall: *r*^2^*/*VE *<* 15% everywhere, indicating that the practical bottleneck is not only information availability (variance coverage) but information exploitation: even within high-variance bases, current models extract only a fraction of the available signal.

---

**Figure ED1:**
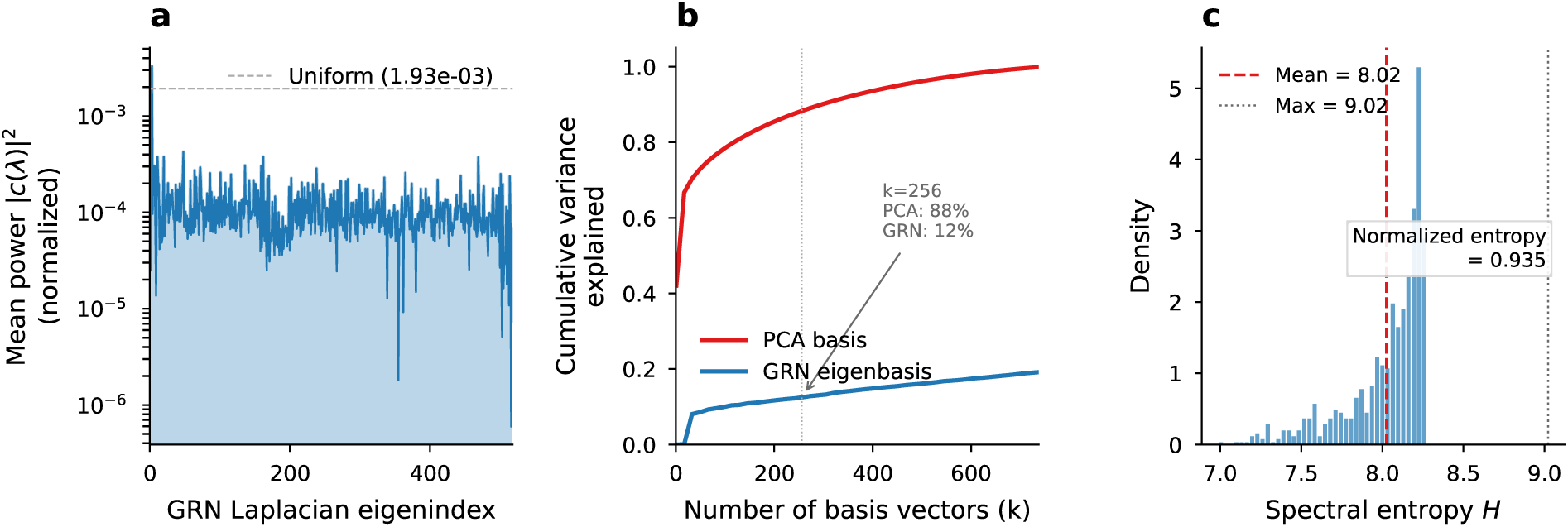
Drug responses are spectrally diffuse on the association graph. **a**, Mean GFT power spectrum |*c*(*λ*)| ^2^ (log scale) across all 751 K562 drug deltas. The nearly flat profile indicates drug signals are distributed across all network Laplacian eigenfrequencies, not concentrated at low frequencies. Dashed line: uniform distribution reference. **b**, Cumulative variance explained by the top-*k* basis vectors. PCA concentrates 88.3% of drug-delta variance in 256 components (7.1 × more than the network eigenbasis at 12.5%). **c**, Per-drug spectral entropy distribution (*n* = 751 drugs). Normalized entropy = 0.935 (close to 1.0, the maximum for a uniform distribution), confirming that no individual drug is spectrally concentrated on the association graph. This spectral diffuseness is consistent across cell lines (A549: 0.933; MCF7: 0.890).

**Table ED1:** Architecture independence: PerturbationMLP (3-layer residual MLP, 525K parameters) reproduces the same two-regime pattern as WPN (19.6M parameters) on K562 sciPlex3. Both models use identical PCA-256 and GRN-256 bases. MLP ± values: standard deviation across 3 random seeds.

| Model | Basis | Params | dir_acc↑ | Pearson↑ | MSE<sub>20</sub>↓ |
| --- | --- | --- | --- | --- | --- |
| PerturbationMLP | GRN-spectral | 525K | 0.418±.004 | 0.004±.010 | 1.121±.020 |
| PerturbationMLP | PCA | 525K | 0.610±.002 | 0.164±.008 | 0.135±.007 |
| WPN | GRN-spectral | 19.6M | 0.433±.029 | 0.029±.049 | 1.100±.037 |
| WPN | PCA | 19.6M | 0.602±.003 | 0.168±.017 | 0.137±.007 |

**Figure ED2:**
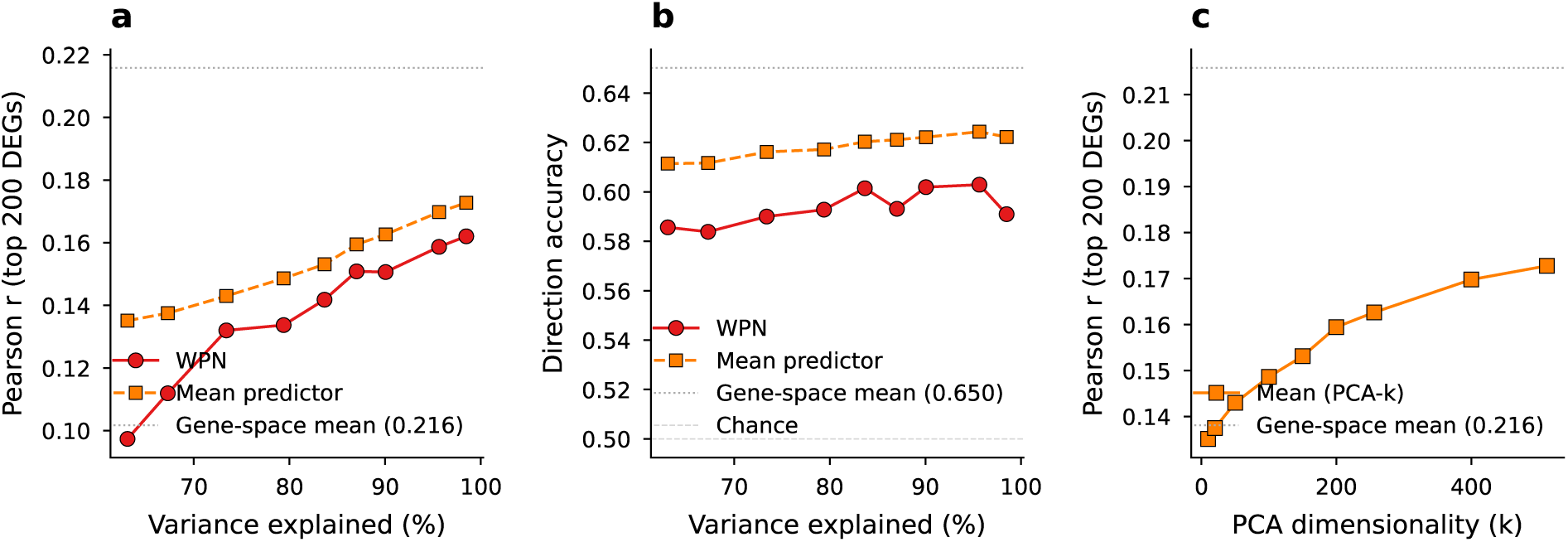
PCA dimensionality sweep reveals monotonic transition. **a**, Pearson *r* (top 200 DEGs) vs. variance explained for WPN (red) and mean predictor (orange) across PCA-*k* bases (*k* = 10–512). Both metrics increase smoothly with variance coverage; the mean predictor outperforms the WPN at every *k*. **b**, Direction accuracy vs. variance explained. Even PCA-10 (63.1% variance) exceeds chance (0.50). **c**, Mean predictor convergence: Pearson *r* at each *k* approaching the gene-space mean (*r* = 0.216, dashed line). At *k* = 512 (98.5% variance), the projected mean reaches *r* = 0.173; the remaining gap reflects the information lost by projection.

**Figure ED3:**
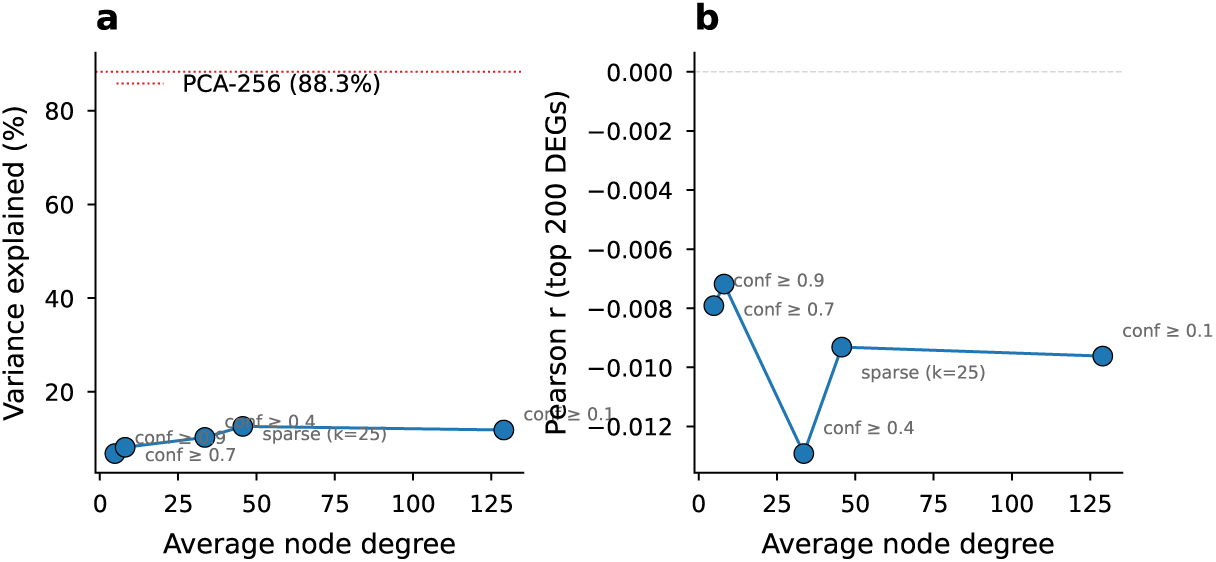
Network sparsity does not rescue the eigenbasis. **a**, Variance explained vs. average node degree for network Laplacian eigenbases at different confidence thresholds (0.1–0.9) and a pre-sparsified variant (*k*=25 edges per node). All configurations remain below 20% variance, far from the PCA-256 reference (88.3%, dashed line). **b**, Pearson *r* vs. average degree. Sparser networks do not improve prediction accuracy.

**Figure ED4:**
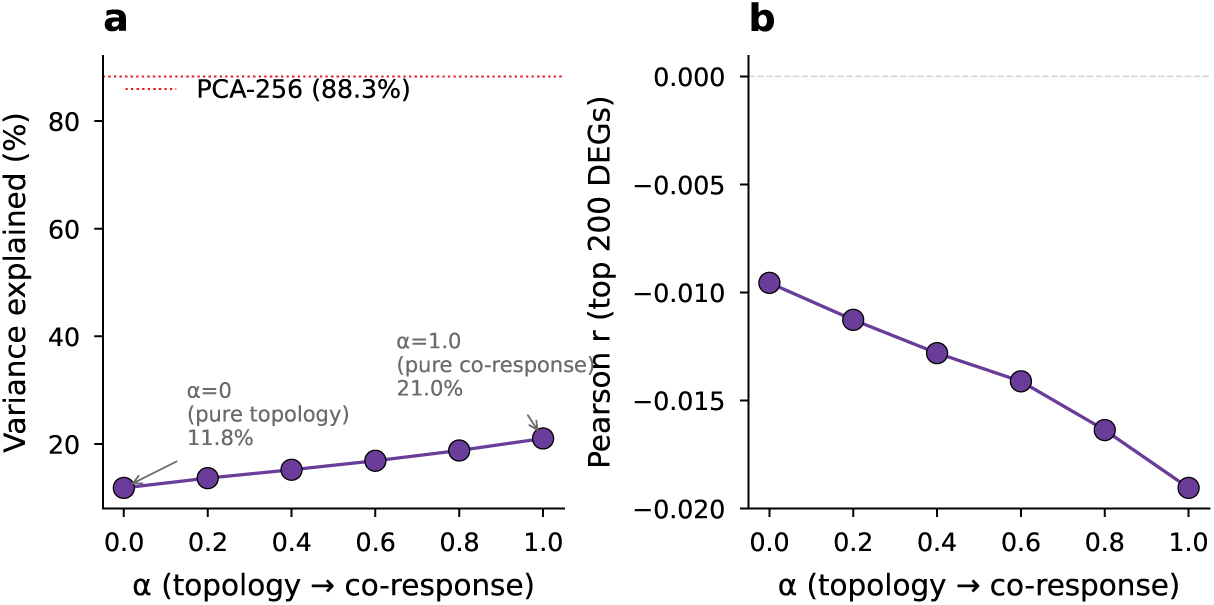
Perturbation-informed reweighting does not reach sufficient variance coverage. **a**, Variance explained vs. *α* (0.0 = pure topology, 1.0 = pure drug co-response). Even at *α*=1.0, the eigenbasis captures far less variance than PCA-256 (dashed line). **b**, Pearson *r* vs. *α*. No value of *α* produces above-chance prediction, demonstrating that the spectral smoothness constraint limits network-based projections.

## Appendix 1: Pooled-control artifact in sciPlex3

sciPlex3 profiles three cell lines (K562, A549, MCF7) under identical drug panels. A natural approach is to pool control cells across cell lines into a single reference, compute drug deltas ***δ*** = ***x̄***_treat_ − ***x̄***_ctrl,pooled_, and train on all conditions jointly. We show that this inflates prediction accuracy by confounding drug effects with cell-line identity.

### Mechanism

K562 (suspension, myeloid), A549 (adherent, lung epithelial), and MCF7 (adherent, breast epithelial) have large basal expression differences. When controls are pooled, the drug delta for a K562-treated condition includes the offset ***x̄***_K562,ctrl_ − ***x̄***_pooled,ctrl_, which reflects cell-line identity, not drug effect. A model can achieve high direction accuracy by simply predicting this cell-line offset for each condition.

### Experiment

We trained WPN on all three cell lines with pooled controls, then re-evaluated using per-cell-line controls (***x̄***_ctrl_ computed separately for each cell line). Performance collapsed to chance (Table S1), confirming that the model had learned cell-line identity rather than drug responses.

**Table S1:** Pooled-control artifact diagnostic. WPN trained with pooled controls, evaluated with per-cell-line controls. All metrics collapse to chance, indicating the model learned cell-line identity, not drug effects.

| Cell line | dir_acc | Pearson | MSE<sub>20</sub> | mag_ratio |
| --- | --- | --- | --- | --- |
| K562 | 0.421 | −0.010 | 1.15 | 3.88 |
| A549 | 0.492 | +0.048 | 6.90 | 3.70 |
| MCF7 | 0.467 | −0.096 | 37.3 | 7.71 |

### Correction

All experiments in the main text use per-cell-line controls. With this correction, GRN-spectral predictions drop to chance (dir_acc = 0.433 ± 0.029, *r* = 0.029 ± 0.049 on K562; 3 seeds), while PCA-based predictions achieve modest but genuine above-chance performance (dir_acc = 0.602 ± 0.003, *r* = 0.168 ± 0.017). The two-regime pattern (GRN at chance, PCA above chance) reproduces across all three cell lines (Table 1).

## Appendix 2: Norman 2019 CRISPRa, dimensionality sweep and mechanism

Test 3 in the main text establishes that on Norman 2019 CRISPRa the mean predictor favours PCA (100% vs. 19.2% VE) while both parameterised architectures invert this ranking and favour the network basis (Table 5). We test a dimensionality sweep to rule out a *k* = *n*_train_ overfitting artefact and the mechanistic analysis of the learned potential landscapes that is summarised in the main text.

### Dimensionality sweep (Tables S2, S3)

We tested *k* ∈ {25, 50, 100, 150, 189} on both WPN and MLP architectures. GRN outperforms PCA at every *k*, including *k* = 25 (where *k/n*_train_ = 0.13 and PCA still captures 87.3% of training-delta variance). PCA–WPN dir_acc decreases monotonically with *k*, consistent with high-dimensional PCA spaces overwhelming a small-sample learner with uninformative directions, while the network’s low-dimensional manifold acts as an implicit regulariser. The MLP (79K–163K parameters) reproduces the WPN pattern almost exactly, ruling out the WPN’s specific potential-gradient inductive bias as the cause of the inversion.

### Why GRN wins for parameterised models on CRISPRa

On the learned potential landscapes at *k* = 189, both architectures converge to effectively rank-1 velocity fields, so the question is which direction each learns. GRN-WPN is substantially better aligned with the true deltas (median cosine similarity = 0.917 vs. 0.785 for PCA-WPN; mean 0.758 vs. 0.668), and its predicted magnitudes are better calibrated (ratio = 1.03 ± 0.71 vs. 0.90 ± 0.64). To distinguish a basis-geometry effect from an easier-optimisation effect, we computed the condition number *κ* of the Hessian of the learned potential *U* at the control state ***s***_ctrl_, defined as the ratio of its largest to smallest eigenvalue; lower *κ* corresponds to a more isotropic, better-conditioned loss surface. PCA-WPN has the lower value (*κ* = 373 vs. 1,013 for GRN-WPN), meaning PCA has the better-conditioned optimisation landscape and yet still loses to GRN; the inversion is therefore a geometry effect, not an optimisation effect. The network basis restricts the model’s gradient direction to a subspace aligned with gene-association structure, which for CRISPRa perturbations (whose effects propagate along pathway edges) provides an informative directional prior. PCA, by contrast, provides *k* roughly equally weighted dimensions with no directional prior. The two-regime pattern for parameterised models is therefore qualitatively distinct from the variance-ceiling governing the mean predictor: availability (VE) transfers across modalities, but exploitation depends on whether the basis geometry aligns with the perturbation’s propagation pattern.

**Table S2:** Norman *k*-sweep: WPN performance as a function of basis dimensionality (*n*_train_ = 189). GRN-WPN outperforms PCA-WPN at every *k*, ruling out *k* = *n*_train_ overfitting as the explanation. PCA-WPN dir_acc decreases with *k*, suggesting that additional PCA dimensions hinder the dynamics model.

| $k$ | WPN-GRN | | | WPN-PCA | | |
| --- | --- | --- | --- | --- | --- | --- |
|  | Var. expl. | dir_acc | Pearson | Var. expl. | dir_acc | Pearson |
| 25 | 10.4% | **0.714** | **0.126** | 87.3% | 0.606 | −0.050 |
| 50 | 11.9% | **0.705** | **0.123** | 93.6% | 0.595 | −0.029 |
| 100 | 14.1% | **0.700** | **0.127** | 97.8% | 0.566 | 0.003 |
| 150 | 17.1% | **0.703** | **0.134** | 99.5% | 0.551 | 0.014 |
| 189 | 19.2% | **0.714** | **0.132** | 100% | 0.579 | 0.010 |

**Table S3:** Norman MLP *k*-sweep: a residual MLP (hidden dim 256, 3 layers, 79K–163K parameters) reproduces the WPN inversion at every *k*, confirming that the network advantage on Norman is architecture-general, not specific to the potential-gradient model.

| $k$ | MLP-GRN | | | MLP-PCA | | |
| --- | --- | --- | --- | --- | --- | --- |
|  | Var. expl. | dir_acc | Pearson | Var. expl. | dir_acc | Pearson |
| 25 | 10.4% | **0.714** | **0.125** | 87.3% | 0.603 | −0.058 |
| 50 | 11.9% | **0.704** | **0.123** | 93.6% | 0.597 | −0.029 |
| 100 | 14.1% | **0.700** | **0.127** | 97.8% | 0.590 | 0.010 |
| 150 | 17.1% | **0.705** | **0.133** | 99.5% | 0.577 | 0.011 |
| 189 | 19.2% | **0.712** | **0.132** | 100% | 0.582 | 0.011 |

**Figure S1:**
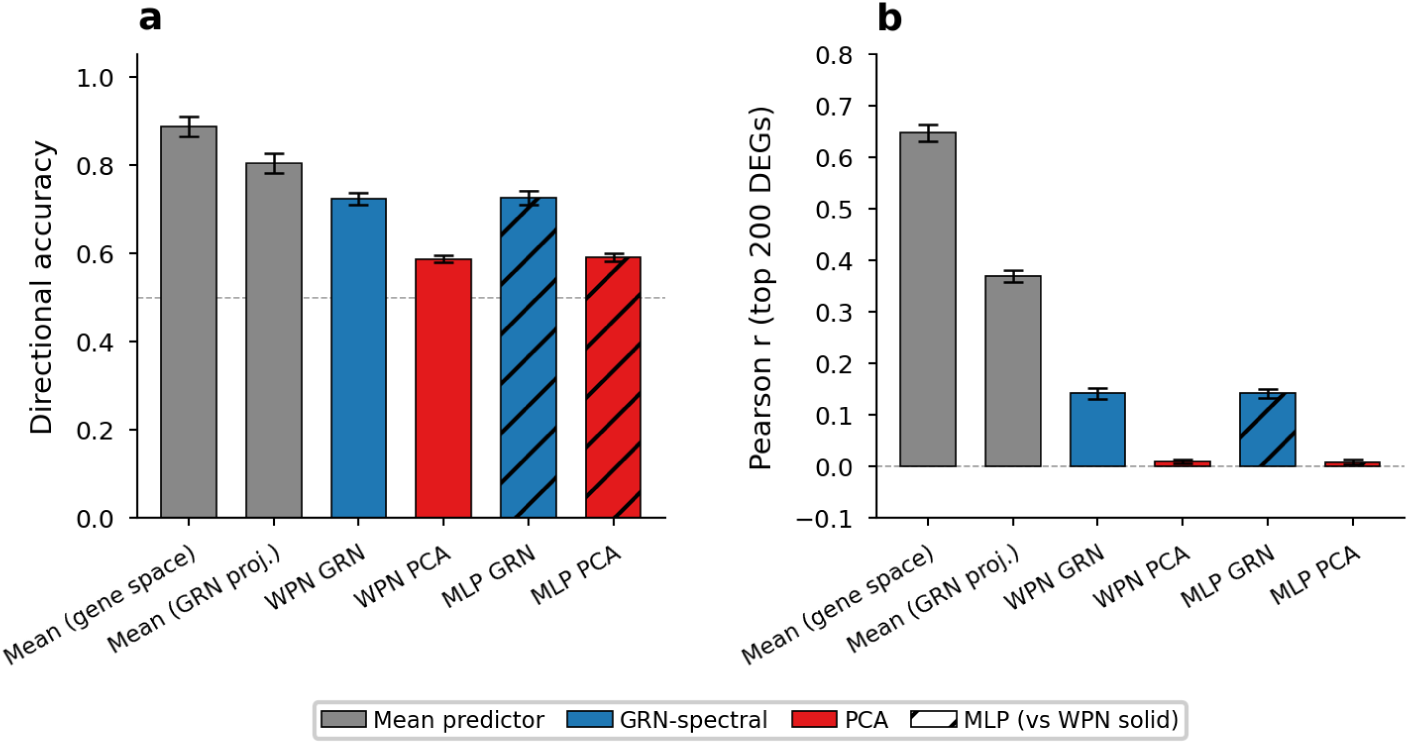
Multi-seed Norman 2019 CRISPRa results (*k* = 189, 3 random seeds). **a**, Directional accuracy and **b**, Pearson *r* (top 200 DEGs) for each method. The mean predictor dominates both learned models; within learned models, the network basis consistently outperforms PCA (the ranking inversion relative to sciPlex3). Error bars: standard deviation across seeds. Hatched bars denote the MLP; solid bars denote the WPN. See Table 5 for numerical values.

**Figure S2:**
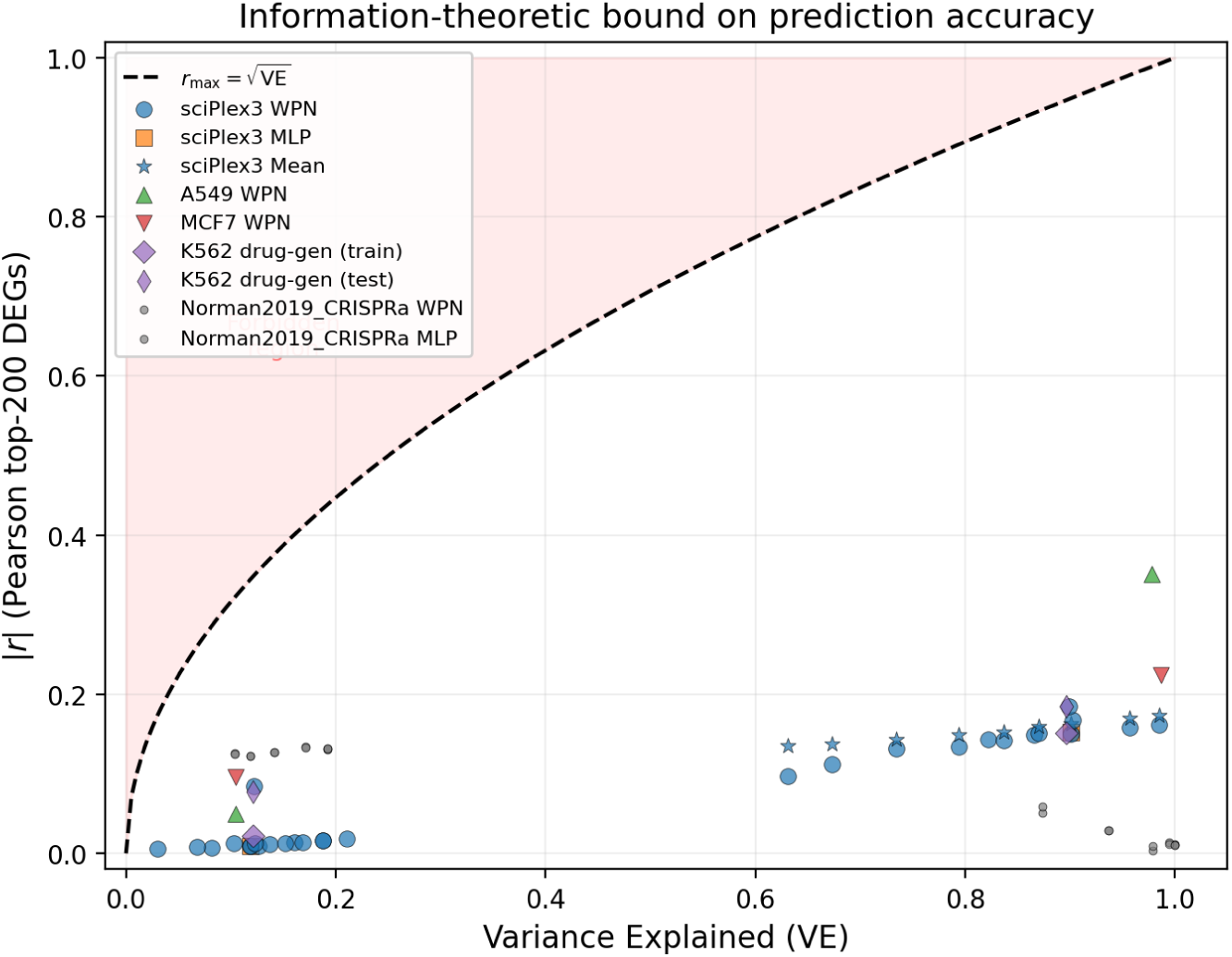
Projection bound on prediction a<u>ccu</u>racy. For any predictor 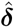 in the range of an orthonormal basis **Φ**, Corr(***δ***, 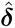)^2^ ≤ VE(**Φ**), i.e., |*r*| ≤ 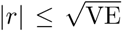 (dashed curve). All 72 data points across both datasets, three architectures (WPN, MLP, mean predictor), and multiple basis types satisfy the bound. The bound itself is a standard consequence of orthogonal projection geometry (Cauchy–Schwarz); the more interesting observation is how far below it all points fall: *r*^2^*/*VE *<* 15% everywhere, indicating that the practical bottleneck is information exploitation, not just information availability.

**Figure S3:**
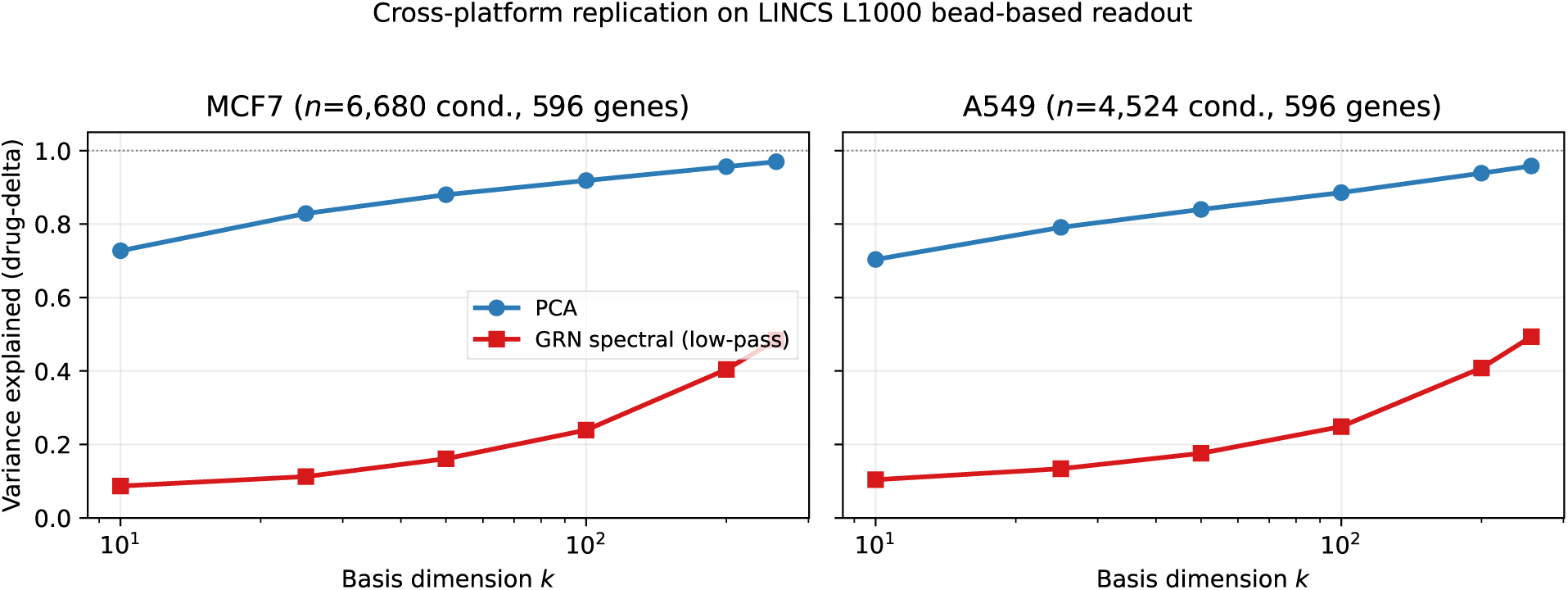
Cross-platform replication on LINCS L1000. PCA versus GRN low-pass spectral basis on the bead-based L1000 platform, across MCF7 (*n* = 6,680 filtered conditions) and A549 (*n* = 4,524 filtered conditions). PCA captures 95–97% of drug-delta variance at *k* = 256 while the GRN low-pass basis remains below 50%, reproducing the sciPlex3 pattern on a completely different measurement modality and cell lines. Data underlying Table 3.

**Figure S4:**
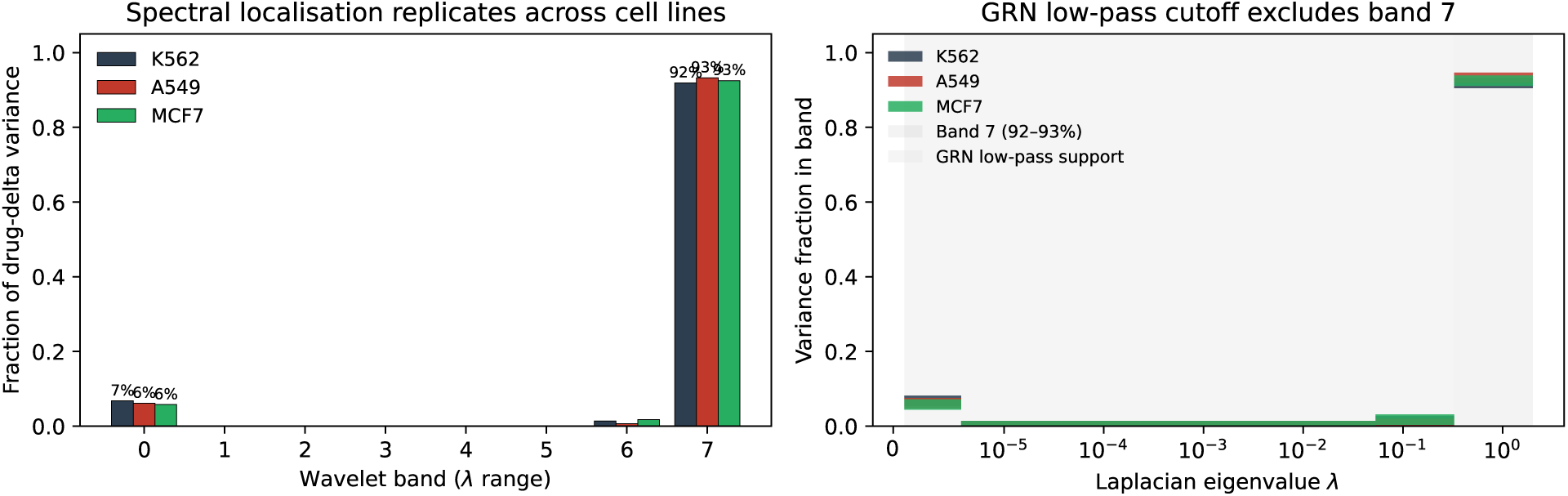
Spectral localisation of drug response replicates across cell lines. **Left:** fraction of drug-delta variance carried by each of eight logarithmic Laplacian frequency bands, for all three sciPlex3 cell lines (*k* = 256). Band 7 (*λ* ∊ [0.326, 2.0], containing 8,235 of 8,288 eigenvectors) accounts for 91.9%, 93.3%, and 92.5% of total drug-delta variance on K562, A549, and MCF7 respectively. **Right:** the same data plotted against the continuous eigenvalue axis, showing that the band sampled by the 256-mode low-pass GRN eigenbasis (light grey, *λ* ≲ 0.33) is almost disjoint from the band carrying the drug-response signal. The GRN basis and the drug signal therefore live in nearly orthogonal subspaces of the same Laplacian, and this spectral mismatch is cell-line-independent.

**Figure S5:**
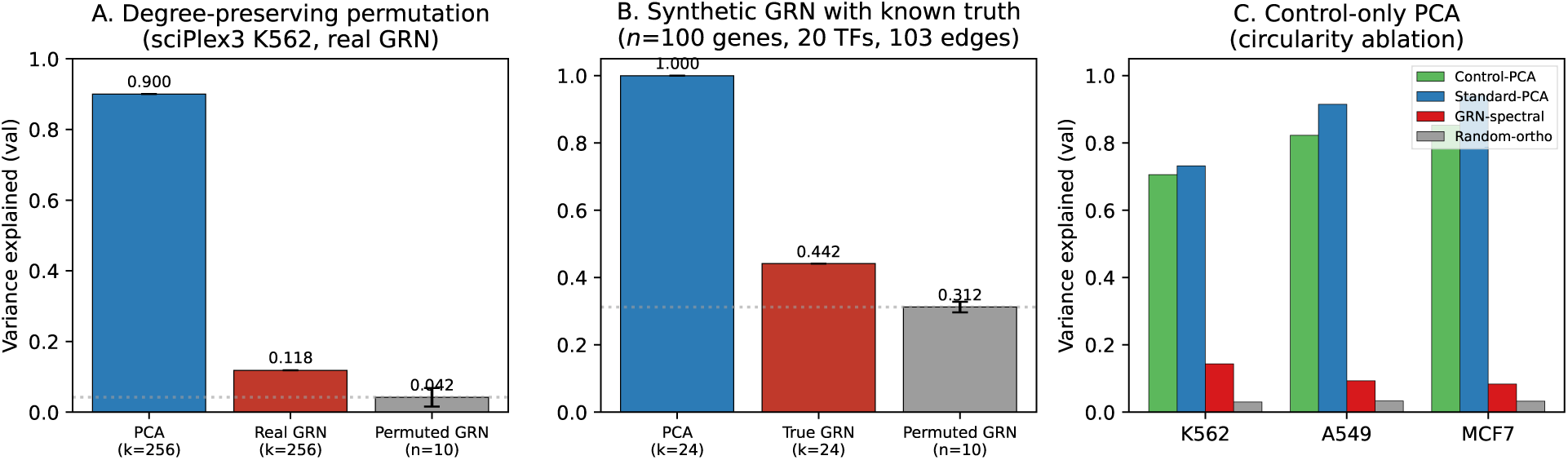
Mechanistic controls for the GRN-basis underperformance. **A** Degree-preserving permutation null on sciPlex3 K562: real GRN (0.118) exceeds the permuted-null mean (0.030 ± 0.002, *n* = 10) with *p <* 10^−4^, while both sit far below PCA (0.900). The basis extracts genuine topology, just not enough of it. **B** Synthetic linear GRN with known ground truth (*n_g_*= 100, 20 TFs, 103 edges, *k* = 24): true GRN (0.442) beats the permuted null (0.312 ± 0.015) confirming the framework detects real structure; PCA still dominates at 1.00. **C** PCA fit on untreated controls alone (“Control-PCA”, green) explains 71–85% of drug-delta variance across K562, A549 and MCF7, rivaling drug-delta-fit Standard-PCA and ruling out circularity. Data underlying Supplementary Tables S11, S14, S15.

**Table S4:** Graph wavelet bases on sciPlex3 (*k* = 256). By using the network’s full frequency spectrum (not just low-frequency modes), band-pass and diffusion wavelets recover nearly all of drug-delta variance, closely matching PCA, while the standard network eigenbasis (low-pass only) captures an order of magnitude less. The pattern replicates across all three sciPlex3 cell lines (K562, A549, MCF7): drug-response signal is distributed across all graph frequencies, and the network topology contains substantially more task-relevant information than the low-pass eigenbasis suggests. A549 and MCF7 PCA/GRN reference MLP performances are reported with the WPN architecture in Table 1.

| Basis | VE | Mean $r$ | MLP $r$ |
| --- | --- | --- | --- |
| K562 |  |  |  |
| PCA (reference) | 89.4% | 0.216 | 0.153 |
| GRN low-pass (reference) | 11.8% | −0.010 | −0.010 |
| Band-pass ($J = 8$) | 88.1% | 0.211 | 0.155 |
| Spectral wavelet ($J = 8$) | 82.0% | 0.211 | 0.093 |
| Diffusion wavelet ($J = 8$) | 88.8% | 0.211 | 0.156 |
| A549 |  |  |  |
| PCA (reference) | 97.5% | 0.328 | — |
| GRN low-pass (reference) | 10.4% | — | — |
| Band-pass ($J = 8$) | 97.2% | 0.328 | 0.320 |
| Spectral wavelet ($J = 8$) | 91.0% | 0.328 | 0.281 |
| Diffusion wavelet ($J = 8$) | 97.1% | 0.328 | 0.322 |
| MCF7 |  |  |  |
| PCA (reference) | 98.5% | 0.236 | — |
| GRN low-pass (reference) | 10.2% | — | — |
| Band-pass ($J = 8$) | 98.4% | 0.236 | 0.228 |
| Spectral wavelet ($J = 8$) | 92.4% | 0.236 | 0.163 |
| Diffusion wavelet ($J = 8$) | 98.3% | 0.236 | 0.230 |

**Table S5:** Variance-explained vs. performance curve on K562 sciPlex3. Bases spanning the full 0–90% VE range, evaluated with a mean predictor and 3-layer residual MLP (40 epochs). Above-chance MLP predictions (dir_acc *>* 0.50) emerge only when VE exceeds ~60%, confirming a monotonic, continuous VE–performance relationship with no threshold or discontinuity. “TopPCA-*k*” takes the top *k* PCA components of the training deltas, filling the previously empty 19–63% VE gap. “RandPCA-*N*” selects *N* PCA columns at random (missing the top components, hence low VE despite using PCA directions). “Hybrid-*m*/*n*” concatenates *m* GRN + *n* PCA dimensions. “Random-ortho-*k*” uses a *k*-dimensional random orthogonal basis.

| Basis | $k$ | VE | Mean $r$ | MLP $\text{dir\_acc}$ | MLP $r$ |
| --- | --- | --- | --- | --- | --- |
| Low variance (<20% VE): all at chance |  |  |  |  |  |
| RandPCA-5 | 5 | 1.4% | 0.021 | 0.401 | −0.007 |
| Random-ortho | 256 | 3.0% | −0.006 | 0.402 | −0.006 |
| Random-ortho-512 | 512 | 6.1% | 0.144 | 0.403 | −0.007 |
| RandPCA-20 | 20 | 9.0% | 0.060 | 0.401 | −0.022 |
| GRN-spectral | 256 | 11.8% | −0.010 | 0.421 | −0.010 |
| Random-ortho-1000 | 1000 | 12.1% | 0.164 | 0.406 | −0.002 |
| Pert-informed (0.8) | 256 | 18.8% | −0.016 | 0.436 | −0.016 |
| Mid variance (50–65% VE): above chance, monotonic |  |  |  |  |  |
| TopPCA-3 | 3 | 54.0% | 0.132 | 0.608 | 0.132 |
| TopPCA-5 | 5 | 58.2% | 0.135 | 0.611 | 0.135 |
| TopPCA-7 | 7 | 60.6% | 0.137 | 0.612 | 0.137 |
| PCA-10 | 10 | 63.1% | 0.097 | 0.586 | 0.097 |
| Gap-filling (65–80% VE): monotonically above chance |  |  |  |  |  |
| Hybrid-240/16 | 256 | 66.8% | 0.138 | 0.589 | 0.108 |
| Hybrid-224/32 | 256 | 70.9% | 0.140 | 0.594 | 0.130 |
| Hybrid-208/48 | 256 | 73.7% | 0.144 | 0.590 | 0.133 |
| Hybrid-192/64 | 256 | 75.8% | 0.147 | 0.599 | 0.140 |
| Hybrid-160/96 | 256 | 79.3% | 0.150 | 0.596 | 0.139 |
| High variance (>80% VE): stable performance |  |  |  |  |  |
| Hybrid-128/128 | 256 | 82.2% | 0.144 | 0.588 | 0.144 |
| PCA-256 | 256 | 90.1% | 0.152 | 0.602 | 0.168 |

**Table S6:** Cell-type-specific network on K562 sciPlex3 (*k* = 256). Gene-gene Pearson correlation from K562 control cells (*n* = 3,935), thresholded at |*r*| ≥ *τ*. The correlation-based network peaks at 19–21% VE (threshold 0.2–0.3), a 1.7 × improvement over the STRING network (12.1%), but still far below PCA (90.0%). Very dense graphs (*τ* = 0.05: avg. degree 1,090) lose specificity; very sparse graphs (*τ* = 0.5: avg. degree 0.36) lose coverage.

| $\tau$ | Edges | Avg. deg. | VE | Mean $r$ |
| --- | --- | --- | --- | --- |
| 0.05 | 4,518K | 1,090 | 8.6% | 0.107 |
| 0.10 | 899K | 217 | 8.9% | 0.062 |
| 0.20 | 62K | 15.1 | 19.3% | 0.012 |
| 0.30 | 12K | 2.9 | 20.7% | 0.073 |
| 0.50 | 1.5K | 0.36 | 2.6% | 0.026 |
| STRING network (ref.) |  | 129 | 12.1% | 0.052 |
| PCA (ref.) |  | — | 90.0% | 0.163 |

**Table S7:** Per-drug-class stratification on K562 sciPlex3 (*k* = 256). Each of the 18 pathway classes shows PCA variance coverage far exceeding GRN, and PCA-projected mean predictor Pearson *r* consistently above GRN-projected mean *r*. No drug class is preferentially GRN-aligned.

| Pathway class | $N$ | $\mathbf{VE}_{\text{PCA}}$ | $\mathbf{VE}_{\text{GRN}}$ | Mean-PCA $r$ | Mean-GRN $r$ |
| --- | --- | --- | --- | --- | --- |
| Cell Cycle | 83 | 91.7% | 15.9% | 0.220 | 0.051 |
| Cytoskeletal Signaling | 48 | 87.0% | 7.2% | 0.387 | 0.093 |
| Epigenetics | 152 | 86.0% | 8.7% | 0.193 | 0.063 |
| Angiogenesis | 44 | 84.1% | 10.3% | 0.299 | 0.061 |
| JAK/STAT | 80 | 84.1% | 11.9% | 0.142 | 0.054 |
| MAPK | 16 | 82.6% | 10.3% | 0.275 | 0.069 |
| Protein Tyrosine Kinase | 64 | 81.8% | 10.9% | 0.182 | 0.063 |
| DNA Damage | 108 | 80.3% | 13.9% | 0.095 | 0.038 |
| Apoptosis | 24 | 78.5% | 8.4% | 0.189 | 0.103 |
| PI3K/Akt/mTOR | 12 | 76.7% | 11.8% | 0.141 | 0.048 |
| TGF-β/Smad | 12 | 76.4% | 6.5% | 0.095 | 0.024 |
| Proteases | 8 | 75.5% | 8.3% | 0.097 | 0.022 |
| Others | 16 | 74.0% | 11.9% | 0.134 | 0.051 |
| Metabolism | 20 | 66.7% | 4.2% | -0.062 | 0.004 |
| Ubiquitin | 8 | 65.2% | 10.4% | 0.021 | 0.038 |
| Neuronal Signaling | 16 | 64.8% | 8.9% | 0.038 | 0.001 |
| Endocrinology | 24 | 63.8% | 8.5% | 0.077 | 0.062 |
| GPCR & G Protein | 8 | 59.3% | 8.2% | 0.070 | 0.050 |

**Table S8:**
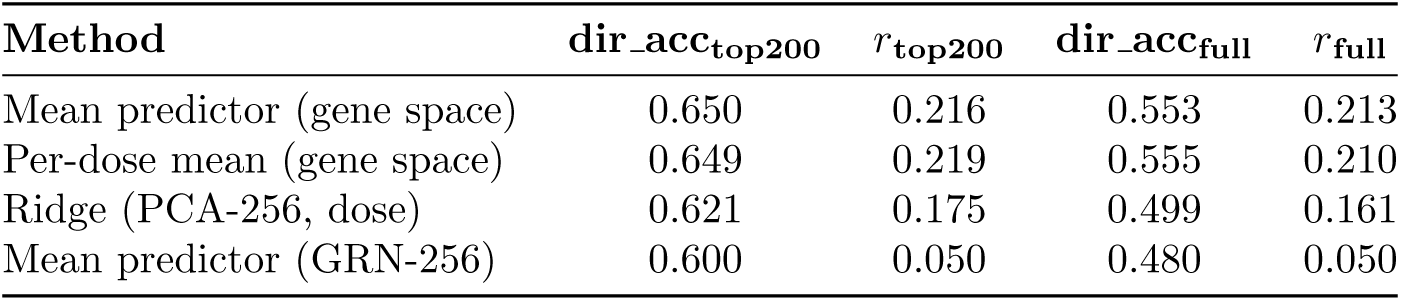
Full-gene metrics robustness check (K562, *k* = 256). Direction accuracy and Pearson *r* computed on all 8,288 genes (excluding near-zero genes with *δ <* 10^−6^) confirm the PCA ≫ GRN gap is not an artifact of top-200 DEG selection.

**Table S9:** PerturBench-style metrics ^24^ for all baselines across three cell lines. Cosine logfc measures cosine similarity between predicted and true log fold-changes. Cosine rank logfc is a pairwise rank metric quantifying per-condition discrimination (0 = perfect, 0.5 = random). All condition-invariant predictors show rank ≈ 0.5, which is the expected outcome under sciPlex3’s shared-control design: models that receive no condition-specific input necessarily produce identical predictions for all conditions. The per-dose mean, which conditions on dose, achieves marginally better rank (0.448 on MCF7), confirming that the metric detects discrimination when condition-specific information is available. WPN and MLP predictions under shared-control input are also condition-invariant and would exhibit the same collapse; baselines are reported here because trained models approximate the mean predictor in this regime.

| Cell line | Method | cosine_logfc↑ | rank_logfc↓ | Pearson↑ | dir_acc↑ |
| --- | --- | --- | --- | --- | --- |
| K562 | Mean predictor (gene) | 0.214 | 0.497 | 0.216 | 0.650 |
|  | Per-dose mean | 0.211 | 0.477 | 0.219 | 0.649 |
|  | Ridge (PCA-256, dose) | 0.164 | 0.483 | 0.175 | 0.621 |
|  | Mean (GRN-256 proj.) | 0.074 | 0.497 | 0.050 | 0.600 |
| A549 | Mean predictor (gene) | 0.276 | 0.497 | 0.346 | 0.654 |
|  | Per-dose mean | 0.275 | 0.483 | 0.344 | 0.648 |
|  | Ridge (PCA-256, dose) | 0.272 | 0.487 | 0.348 | 0.643 |
|  | Mean (GRN-256 proj.) | 0.046 | 0.497 | 0.176 | 0.548 |
| MCF7 | Mean predictor (gene) | 0.206 | 0.497 | 0.241 | 0.616 |
|  | Per-dose mean | 0.227 | 0.448 | 0.277 | 0.622 |
|  | Ridge (PCA-256, dose) | 0.189 | 0.484 | 0.243 | 0.597 |
|  | Mean (GRN-256 proj.) | 0.043 | 0.497 | 0.091 | 0.563 |

**Table S10:** scGen-style VAE basis comparison on K562 sciPlex3 (*k* = 256). A variational autoencoder trained with latent-space arithmetic (following Lotfollahi et al. ^3^) reproduces the PCA ≫ GRN basis effect observed with the WPN and MLP. scGen-PCA matches WPN direction accuracy (0.60 vs. 0.60); scGen-GRN collapses to below chance (0.45), consistent with the information-loss mechanism. scGen-PCA achieves high direction accuracy but strikingly low Pearson *r* (0.029 vs. 0.168 for WPN-PCA at comparable dir_acc). This dissociation likely reflects VAE reconstruction noise: the latent-space arithmetic recovers the correct sign of gene changes (hence high dir_acc) but introduces magnitude distortions through the decoder that degrade Pearson correlation, which is magnitude-sensitive. The elevated MSE_20_ (0.536 vs. 0.134 for WPN-PCA) supports this interpretation: the VAE’s reconstructed predictions have correct direction but poorly calibrated magnitudes. This is consistent with the broader finding that perturbation prediction models struggle to exceed linear baselines ^4,5^.

| Model (basis) | dir_acc | $r_{\text{top200}}$ | MSE<sub>top20</sub> | Params |
| --- | --- | --- | --- | --- |
| scGen-VAE (PCA-256) | 0.598 | 0.029 | 0.536 | 364K |
| scGen-VAE (GRN-256) | 0.450 | −0.012 | 1.017 | 364K |
| Reference (from other experiments): |  |  |  |  |
| Mean predictor (PCA) | 0.650 | 0.216 | 0.134 | 0 |
| WPN (PCA-256) | 0.601 | 0.152 | 0.136 | 19.6M |
| MLP (PCA-256) | 0.602 | 0.168 | 0.134 | 525K |
| WPN (GRN-256) | 0.430 | −0.010 | 0.147 | 19.6M |

## Appendix 3: Causal tests of network topology

In this work we establish that the gene-gene association network eigenbasis captures only 11.8% of drug-response variance on sciPlex3, compared to 90% for PCA. Two questions follow naturally. First, does the specific biological topology of the association network contribute to that 11.8%, or would any network with the same degree distribution perform equally? Second, does the failure reflect the use of an undirected association graph rather than a true directed regulatory network? We designed four complementary tests to address these questions, ranging from statistical null models on real data to a fully synthetic positive control.

### Control A: Degree-preserving permutation null

To test whether biological network topology contributes information beyond what is implied by the degree distribution alone, we compared the real association network against degree-preserving random permutations. We applied a configuration-model edge-swap algorithm ^21^ that repeatedly selects two random edges and rewires their endpoints, rejecting swaps that would create self-loops or multi-edges. Each permuted network preserves the exact degree sequence of the original graph but destroys the specific wiring pattern that encodes biological relationships. We generated 10 independent permutations (each with 10 × *E* swap attempts; swap acceptance rate ∼77%) and evaluated each as a projection basis on sciPlex3 K562.

The real network eigenbasis captures significantly more variance than the permuted null (VE = 11.8% vs. 4.2% ± 2.6%; *p <* 10^−4^ by one-sided permutation test; Table S11). Direction accuracy follows the same pattern (0.601 vs. 0.544 ± 0.017; *p <* 10^-4^). The biological wiring of the association network therefore contributes genuine, topology-specific information that degree-matched random graphs do not replicate. However, this advantage remains modest in absolute terms: the real network captures roughly three times the variance of permuted networks, but both fall far below PCA (90.0%).

**Table S11:**
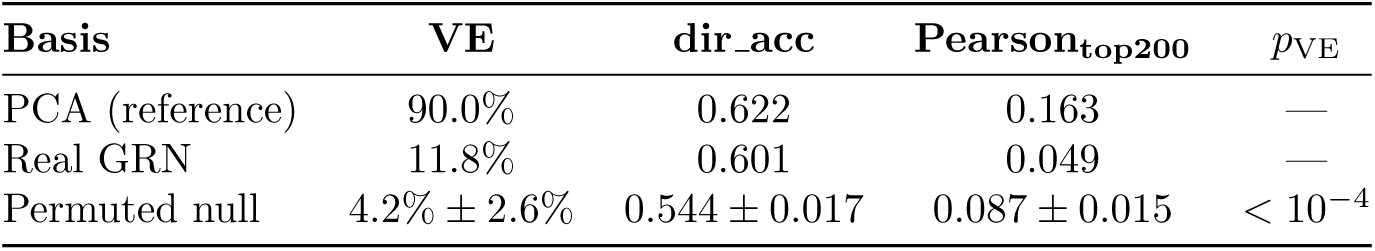
Degree-preserving permutation test on sciPlex3 K562 (*k* = 256, *n* = 10 permutations). The real GRN significantly outperforms degree-matched permuted networks (*p <* 10^−4^), confirming that biological topology contributes information beyond the degree distribution. Both remain far below PCA.

| Basis | VE | dir_acc | Pearson<sub>top200</sub> | $p_{VE}$ |
| --- | --- | --- | --- | --- |
| PCA (reference) | 90.0% | 0.622 | 0.163 | — |
| Real GRN | 11.8% | 0.601 | 0.049 | — |
| Permuted null | $4.2\% \pm 2.6\%$ | $0.544 \pm 0.017$ | $0.087 \pm 0.015$ | $< 10^{-4}$ |

### Control B: Directed regulatory network comparison

The association network used in the main text is primarily undirected: its edges reflect protein-protein interaction, co-expression, and pathway co-membership rather than directed transcription-factor-to-target regulation. To test whether a curated, directed regulatory network provides a better spectral basis, we evaluated two established TF-target databases: CollecTRI ^17^, which aggregates literature-curated signed regulatory interactions, and DoRothEA ^18^, which assigns confidence levels (A through E) to TF-target pairs based on multiple evidence sources.

We computed eigenbases from each network under three spectral decomposition strategies. For CollecTRI, we tested both a standard symmetrized normalized Laplacian (which discards edge directionality) and a signed Laplacian (**L**_signed_ = **D**_|·|_ − (**A**_+_ − **A**_−_), where **A**_+_ and **A**_−_ encode activation and repression edges separately) that preserves the sign structure of regulatory interactions. For DoRothEA, we tested high-confidence (ABC) and broad (ABCDE) subsets using the standard symmetrized Laplacian.

All directed networks substantially underperform the undirected association network (Table S12). The primary bottleneck is coverage: CollecTRI covers only 35.3% of the 8,288-gene vocabulary (14,307 edges from 574 TFs), and DoRothEA ABC covers 46.9% (10,789 edges from 217 TFs), compared to the association network’s 534,670 edges spanning nearly all genes. The resulting spectral bases are constructed from far sparser graphs, limiting the variance they can capture. The signed Laplacian, which preserves activation and repression structure, provides no meaningful advantage over symmetrization (VE = 6.2% vs. 4.7%), suggesting that regulatory sign information does not rescue the sparsity limitation at the spectral level. DoRothEA ABC and ABCDE produce identical results, indicating that the additional lower-confidence edges collapse to the same set at the gene-vocabulary intersection.

**Table S12:** Directed regulatory network comparison on sciPlex3 K562 (*k* = 256). Curated TF-target networks (CollecTRI, DoRothEA) underperform the undirected association network. The primary limitation is coverage: directed databases provide 10–30× fewer edges than the association graph. The signed Laplacian does not rescue this deficit.

| Network | Edges | Coverage | VE | dir_acc | Pearson<sub>top200</sub> |
| --- | --- | --- | --- | --- | --- |
| PCA (reference) | — | — | 90.0% | 0.622 | 0.163 |
| STRING/OmniPath (assoc.) | 534,670 | $\sim 100\%$ | 11.8% | 0.601 | 0.049 |
| CollecTRI (symmetrized) | 14,307 | 35.3% | 4.7% | 0.302 | 0.070 |
| CollecTRI (signed) | 14,307 | 35.3% | 6.2% | 0.304 | 0.066 |
| DoRothEA ABC | 10,789 | 46.9% | 6.1% | 0.533 | 0.057 |
| DoRothEA ABCDE | 10,789 | 46.9% | 6.1% | 0.533 | 0.057 |

### Control C: CRISPRi vs. CRISPRa dissociation

If the association network encodes directional regulatory information, it should differentially predict gene activation (CRISPRa) versus gene repression (CRISPRi): a directed network “knows” which direction perturbation effects propagate, whereas an undirected network treats activation and repression identically. To test this, we compared PCA and GRN eigenbases on two genetic perturbation datasets in the same cell line (K562): Norman et al. 2019^13^ (CRISPRa, 236 single-gene activations) and Replogle et al. 2022^28^ (CRISPRi, 500 single-gene repressions).

The GRN underperforms PCA by a similar margin on both perturbation types (Table S13). The PCA-to-GRN variance gap is 80.8 percentage points on CRISPRa and 75.6 on CRISPRi, a difference of only 5.2 percentage points. Direction accuracy shows a similarly small dissociation (6.7 percentage points). These results are inconsistent with the hypothesis that the association network carries directional regulatory information that preferentially aids one perturbation modality over the other.

We note two important caveats. First, the Norman and Replogle datasets share only 4 overlapping target genes, making this a between-dataset comparison with potential confounds from dataset-specific technical variation. Second, the CRISPRi perturbation effects in Replogle are substantially weaker than the CRISPRa effects in Norman (mean delta norm 1.42 vs. 4.43), which may limit the sensitivity of the comparison. A more definitive test would require a single dataset with matched CRISPRa and CRISPRi perturbations targeting the same genes; to our knowledge, no such dataset is publicly available at sufficient scale.

**Table S13:** CRISPRi vs. CRISPRa dissociation test (K562). The GRN underperforms PCA by a similar margin on both gene activation (CRISPRa) and repression (CRISPRi), indicating that the association network does not encode directional information that preferentially aids either perturbation modality.

| Pert. type | Basis | VE | dir_acc | Pearson<sub>top200</sub> | VE gap |
| --- | --- | --- | --- | --- | --- |
| CRISPRa<br>(Norman, $n = 236$) | PCA ($k = 189$) | 100.0% | 0.868 | 0.642 | 80.8 pp |
| | GRN ($k = 189$) | 19.2% | 0.788 | 0.361 | |
| CRISPRi<br>(Replogle, $n = 500$) | PCA ($k = 256$) | 86.3% | 0.521 | 0.077 | 75.6 pp |
| | GRN ($k = 256$) | 10.7% | 0.508 | 0.055 | |

### Control D: Synthetic ground-truth GRN

The preceding three tests use real biological networks whose true causal structure is unknown. As a positive control, we generated a fully synthetic dataset in which the ground-truth regulatory network is known by construction, allowing us to ask: when the data are genuinely generated by a regulatory network, does the permutation test detect it?

We constructed a scale-free-like regulatory network (100 genes, 20 transcription factors, 103 directed edges) and simulated steady-state gene expression using coupled stochastic differential equations with Hill-function activation dynamics. We generated 500 control cells and 30 single-gene knockout perturbations (200 cells each), then computed eigenbases from the true network, 10 degree-preserving permutations, and 10 random networks with matched density.

The true GRN significantly outperforms both permuted and random networks (Table S14; *p <* 10^−4^ for both comparisons). The VE effect sizes are substantial: the true network captures 44.2% of variance, compared to 31.2% ± 1.5% for permuted and 29.3% ± 2.2% for random networks (effect sizes of +13.0 and +14.9 percentage points, respectively). This confirms that the permutation test framework is sensitive enough to detect genuine network structure when the data are truly generated by a regulatory topology.

However, even with perfect ground-truth knowledge of the generative network, PCA captures ∼100% of variance while the true GRN captures only 44%. This reinforces the main text finding: the spectral smoothness constraint inherent to Laplacian eigenvectors limits variance coverage regardless of how biologically accurate the network is. Perturbation effects (even those generated by a regulatory network) do not propagate exclusively along smooth, low-frequency modes of that network.

**Table S14:** Synthetic ground-truth GRN test (100 genes, 20 TFs, 103 edges, 30 perturbations, *k* = 24). The true network significantly outperforms permuted and random networks, validating the permutation test framework. PCA still dominates even when the generative network is known exactly, confirming that the low-pass spectral constraint is the fundamental limitation.

| Basis | VE | dir_acc | $p_{VE}$ |
| --- | --- | --- | --- |
| PCA | 100.0% | 0.547 | — |
| True GRN | 44.2% | 0.465 | — |
| Permuted GRN ($n = 10$) | $31.2\% \pm 1.5\%$ | $0.395 \pm 0.022$ | $< 10^{-4}$ |
| Random GRN ($n = 10$) | $29.3\% \pm 2.2\%$ | $0.396 \pm 0.029$ | $< 10^{-4}$ |

### Summary

These four controls converge on three conclusions. First, biological network topology contributes real, statistically significant information to the eigenbasis (Controls A and D), ruling out the possibility that the association network eigenbasis is interchangeable with any degree-matched random graph. Second, curated directed regulatory networks do not improve on the undirected association graph (Control B), primarily because current databases are too sparse for genome-scale spectral analysis, and the association network carries no detectable directional information (Control C). Third, even when the true generative network is known perfectly (Control D), the Laplacian eigenbasis captures less than half the variance that PCA achieves, confirming that the low-pass spectral constraint is the binding limitation rather than network accuracy.

## Appendix 4: Basis circularity controls

A potential concern with the PCA-vs-network comparison is that PCA is fit on training drug conditions and then evaluated by variance explained on those same conditions, making PCA the basis “mathematically defined to maximize the exact signal you are evaluating against.” We address this concern with two controls that break the data-sharing link between basis construction and evaluation.

### Control 1: PCA on untreated cells

We computed PCA on the raw control-cell expression matrix (cells with zero drug exposure) for each cell line independently and measured variance explained on drug deltas (Supplementary Table S15). This “control-PCA” basis has never seen any drug-response data, yet it captures 65–85% of drug-response variance across all three cell lines, compared to 10–12% for the network eigenbasis. Mean-predictor performance is virtually identical between control-PCA and standard PCA (Pearson *r* = 0.163 vs. 0.163 on K562; 0.342 vs. 0.343 on A549; 0.236 vs. 0.237 on MCF7), confirming that drug data contributes negligible additional prediction power beyond the basal gene covariance structure.

This result is consistent with the spectral diffuseness finding in the main text: drug responses are distributed across all graph frequencies but are well represented in the low-rank subspace defined by gene covariance in the absence of perturbation. The relevant geometry is not drug-specific; it reflects the cell type’s intrinsic expression manifold.

**Table S15:** Control-PCA ablation. “Control-PCA” is fit on untreated cells only (no drug data). “Standard-PCA” is fit on control + treated expression means (the method used throughout the main text). “Delta-PCA” is fit directly on drug deltas (the most circular case). All bases are 256-dimensional. VE is measured on training-split drug deltas; mean-predictor metrics are computed on the validation split.

| Cell line | Basis | Drug data? | VE↑ | dir_acc↑ | Pearson↑ | MSE<sub>20</sub>↓ |
| --- | --- | --- | --- | --- | --- | --- |
| K562 | Control-PCA | No | 65.1% | 0.619 | 0.163 | 0.135 |
|  | Standard-PCA | Yes | 89.9% | 0.622 | 0.163 | 0.135 |
|  | Delta-PCA | Yes | 89.9% | 0.622 | 0.162 | 0.135 |
|  | GRN-spectral | No | 11.8% | 0.601 | 0.049 | 0.143 |
|  | Random-ortho | No | 3.0% | 0.561 | 0.106 | 0.143 |
| A549 | Control-PCA | No | 83.3% | 0.635 | 0.342 | 0.903 |
|  | Standard-PCA | Yes | 97.7% | 0.640 | 0.343 | 0.899 |
|  | Delta-PCA | Yes | 97.7% | 0.637 | 0.343 | 0.899 |
|  | GRN-spectral | No | 10.5% | 0.546 | 0.175 | 1.009 |
|  | Random-ortho | No | 3.3% | 0.574 | 0.248 | 1.010 |
| MCF7 | Control-PCA | No | 84.5% | 0.595 | 0.236 | 2.169 |
|  | Standard-PCA | Yes | 98.6% | 0.599 | 0.237 | 2.159 |
|  | Delta-PCA | Yes | 98.6% | 0.597 | 0.237 | 2.159 |
|  | GRN-spectral | No | 10.4% | 0.566 | 0.091 | 2.342 |
|  | Random-ortho | No | 3.2% | 0.537 | 0.174 | 2.340 |

### Control 2: Cross-cell-line PCA transfer

To further decouple basis construction from evaluation data, we fit PCA on all drug deltas from one cell line and evaluated variance explained on a different cell line’s drug deltas (Supplementary Table S16). All six transfer directions produce VE far exceeding the network eigenbasis: transferred PCA captures 39–68% of target-cell-line drug-response variance, compared to 10–12% for the GRN.

Transfers into K562 (a suspension myeloid line) from the two adherent epithelial lines (A549, MCF7) show lower VE (∼39%), while transfers between A549 and MCF7 are stronger (54–68%), consistent with the two epithelial lines sharing more gene covariance structure. This asymmetry reflects genuine biological differences in expression geometry rather than any property of the evaluation metric. Even in the weakest transfer direction (A549 → K562, VE = 38.8%), the transferred basis captures 3.3× more drug-response variance than the GRN eigenbasis.

**Table S16:** Cross-cell-line PCA transfer. PCA is fit on all drug deltas from the source cell line and evaluated on the target cell line’s drug deltas (VE on training-split deltas). “Same-cell PCA” uses all deltas from the target cell line (oracle upper bound). All bases are 256-dimensional.

| Transfer | Transferred VE↑ | Same-cell VE | GRN VE | Random VE |
| --- | --- | --- | --- | --- |
| K562 → A549 | 57.7% | 97.2% | 10.5% | 3.3% |
| K562 → MCF7 | 64.3% | 98.4% | 10.4% | 3.2% |
| A549 → K562 | 38.8% | 87.8% | 11.8% | 3.0% |
| A549 → MCF7 | 68.2% | 98.4% | 10.4% | 3.2% |
| MCF7 → K562 | 39.5% | 87.8% | 11.8% | 3.0% |
| MCF7 → A549 | 54.1% | 97.2% | 10.5% | 3.3% |

### Interpretation

Together, these controls establish that the PCA-vs-network variance gap is not an artifact of fitting PCA on the same data used for evaluation. PCA bases that have never seen drug data (Control 1) or that were fit on a different cell line’s drug data entirely (Control 2) both capture several-fold more drug-response variance than the network eigenbasis. The finding that PCA outperforms the GRN eigenbasis reflects the biological fact that drug responses are spectrally diffuse on the association graph (main text, Graph Fourier analysis) and are well represented in the low-rank gene covariance subspace that PCA captures. The cell type’s basal expression geometry, not any drug-specific signal, largely determines this subspace: control-PCA achieves nearly identical mean-predictor performance to standard PCA despite having zero access to drug-response data.

